# Analysis of Rod/Cone Gap Junctions from the Reconstruction of Mouse Photoreceptor Terminals

**DOI:** 10.1101/2021.09.06.459091

**Authors:** Munenori Ishibashi, Joyce Keung, Catherine W. Morgans, Sue A. Aicher, James R. Carroll, Joshua H. Singer, Li Jia, Wei Li, Iris Fahrenfort, Christophe P. Ribelayga, Stephen C. Massey

## Abstract

Using serial blockface-scanning electron microscopy (SBF-SEM) and focused ion beam-scanning electron microscopy (FIB-SEM), combined with confocal microscopy for the gap junction protein Cx36, we reconstructed mouse photoreceptor terminals and located the gap junctions between them. An exuberant spray of fine telodendria extends from each cone pedicle (including blue cones) to contact 40-50 nearby rod spherules where Cx36 clusters were located, close to the mouth of the synaptic opening. There were approximately 50 Cx36 clusters per cone pedicle and 2-3 per rod spherule. We were unable to detect rod/rod or cone/cone coupling. Thus, rod/cone coupling accounts for nearly all gap junctions between photoreceptors. Our calculations suggest a mean of 82 Cx36 channels between a rod/cone pair of which 25% are open at rest. Rod/cone gap junctions are modulated by dopamine. Comparing our morphological calculations of maximum coupling to previous physiological results suggests that dopamine antagonists can drive rod/cone gap junctions to a surprisingly high open probability, approaching 100%.

## Introduction

In the mammalian retina, with few exceptions, rods far outnumber cones. Specifically, for the mouse retina, only 3% of the photoreceptors are cones (Carter Dawson and Lavail, 1979). The synaptic endings of photoreceptors, known as cone pedicles and rod spherules, terminate in the outer plexiform layer (OPL) where they make synaptic connections with second-order neurons, horizontal cells and bipolar cells. In general, the retina is a layered structure and the OPL is no exception. Cone pedicles are found in a single layer in the mid-OPL whereas the smaller rod spherules accumulate in several layers between the cone pedicles and the photoreceptor cell bodies in the top (distal) half of the OPL. Numerically, the OPL is dominated by rod spherules.

Cone pedicles are the largest structures in the OPL. They contain numerous synaptic ribbons and make contacts with horizontal cells and 13 types of cone bipolar cell. One of these bipolar cell types, cb9, contacts short wavelength-sensitive (i.e. “blue”) cones selectively, which can be identified on this basis (Behrens et al., 2016; Nadal-Nicolás et al., 2020). Rod spherules contain a single large ribbon and make synapses with the axon terminals of horizontal cells and with the dendrites of a single type of rod bipolar cell. Each rod bipolar cell contacts ∼35 rods (Behrens et al., 2016), forming the most sensitive or primary rod pathway.

In addition to chemical synapses, there are a large number of small gap junctions in the OPL, which were first described in early ultrastructural studies (Kolb, 1977; Raviola and Gilula, 1973; Smith et al., 1986). Gap junctions mediate electrical coupling between neurons and contain clusters of intercellular channels that allow the passage of ions and other small molecules. At a gap junction, the membranes of two neighboring cells are apposed or “zippered”, leaving only a 2-4 nm “gap” (Bloomfield and Völgyi, 2009; Miller and Pereda, 2017). Each gap junction channel is formed of two docked hemichannels, or connexons and each connexon is assembled from six connexin subunits. Connexin expression is required on both sides to form a gap junction (Miller et al.,2017; Jin et al., 2020). The vertebrate connexin family in mouse includes 20 members, including Cx36, the most common neuronal connexin (Beyer and Berthoud, 2009; Söhl et al., 2004). Gap junctions are widespread in neural tissue and support functions such as signal averaging, noise reduction and synchronization (Connors, 2017; Nagy et al., 2018).

In lower vertebrates, rods are extensively electrically coupled (Zhang and Wu, 2004) but, in general, this is not the case for mammals. Cone/cone and rod/cone gap junctions have been reported in cat and primate retinae (Kolb, 1977; Smith et al., 1986). Specifically, basal processes, called telodendria, spread laterally from cone pedicles to make small gap junctions with rod spherules. In freeze-fracture electron microscopy (EM), they appear as strings of single particles curved around the synaptic invagination of rod spherules (Raviola and Gilula, 1973).

The presence of rod/cone gap junctions at rod/cone contacts is consistent with the transmission of rod signals to cones and cone-driven second order neurons such as horizontal cells and cone bipolar cells (Asteriti et al., 2014; Ingram et al., 2019; Li et al., 2010; Nelson, 1977; Ribelayga et al., 2008; Ribelayga and Mangel, 2010; Schneeweis and Schnapf, 1995). More rarely, rod/rod gap junctions have been suggested from tracer coupling studies and from EM studies of the mouse retina (Jin et al., 2015; Li et al., 2012; Tsukamoto et al., 2001).

Rods and cones both express Cx36, but no other connexins (Bloomfield and Völgyi, 2009; Jin et al., 2020; O’Brien et al., 2012). Connexin expression is required on both sides of a gap junction and, in fact, Cx36 mediated rod/cone coupling accounts for most of the gap junctions in the OPL (Asteriti et al., 2017; Ingram et al., 2019; Jin et al., 2020; Miller et al., 2017). If indeed rods express Cx36, then it raises the possibility that they could also be coupled via Cx36 gap junctions. However, the evidence for rod/rod gap junctions in the mammalian retina is mixed (Bloomfield and Völgyi, 2009; Jin et al., 2020; Tsukamoto et al., 2001) and will be addressed here. We used a combination of confocal microscopy, serial blockface scanning electron microscopy (SBF-SEM), and focused-ion beam SEM (FIB-SEM) to address four major goals: i) to reconstruct the cone telodendrial network; ii) to identify all gap junction types in the OPL; iii) to resolve the question of rod/rod coupling, and iv) to make a quantitative estimate of rod/cone coupling.

Gap junctions can be difficult to identify reliably in EM material (Pallotto et al., 2015). For this project, we chose to use a publicly available dataset, e2006 (Helmstaedter et al., 2013), which contains photoreceptor terminals and the OPL, to map membrane contacts between photoreceptors. Unfortunately, because this dataset was prepared to enhance membrane contrast and facilitate tracing neuronal processes, gap junctions are not visible. To work around this problem, we used confocal microscopy to identify the location of Cx36 gap junctions around the synaptic opening of each rod spherule (Jin et al, 2020). Tracing the contacts between cone telodendria and rod spherules revealed numerous potential gap junctions coincident with the location of Cx36. Thus, the combination of confocal microscopy and SBF-SEM is complementary. To confirm the location of rod/cone gap junctions, we used FIB-SEM on new samples processed to preserve the ultrastructure, which allowed high-resolution imaging of contacts between cone pedicles and rod spherules. In addition, FIB-SEM provides isotropic data (same resolution in each dimension), facilitating 3D reconstruction (Xu et al., 2017) and the measurement of gap junction dimensions.

## Results

### Confocal microscopy: Localization of Cx36 in the OPL

Immunofluorescent labeling for Cx36 revealed small clusters of labeling in the OPL (Fig. 1A, red). To locate these potential gap junctions, we used an antibody to the vesicular glutamate transporter (vGlut1) (Fig. 1A, blue) to identify rod spherules and an antibody to cone arrestin to label cone pedicles (Fig. 1A, green). Labeling with the cone arrestin (CAR) antibody stains cones in their entirety from the outer segment to the pedicle (Fig. 1A). Fine processes, called telodendria, extend laterally from each cone pedicle to form an overlapping matrix in the outer plexiform layer (OPL). The area between and above the cone pedicles is filled with numerous vGlut1-labeled rod spherules while the space underneath is occupied by the dendrites of horizontal cells and bipolar cells.

**Figure 1.**
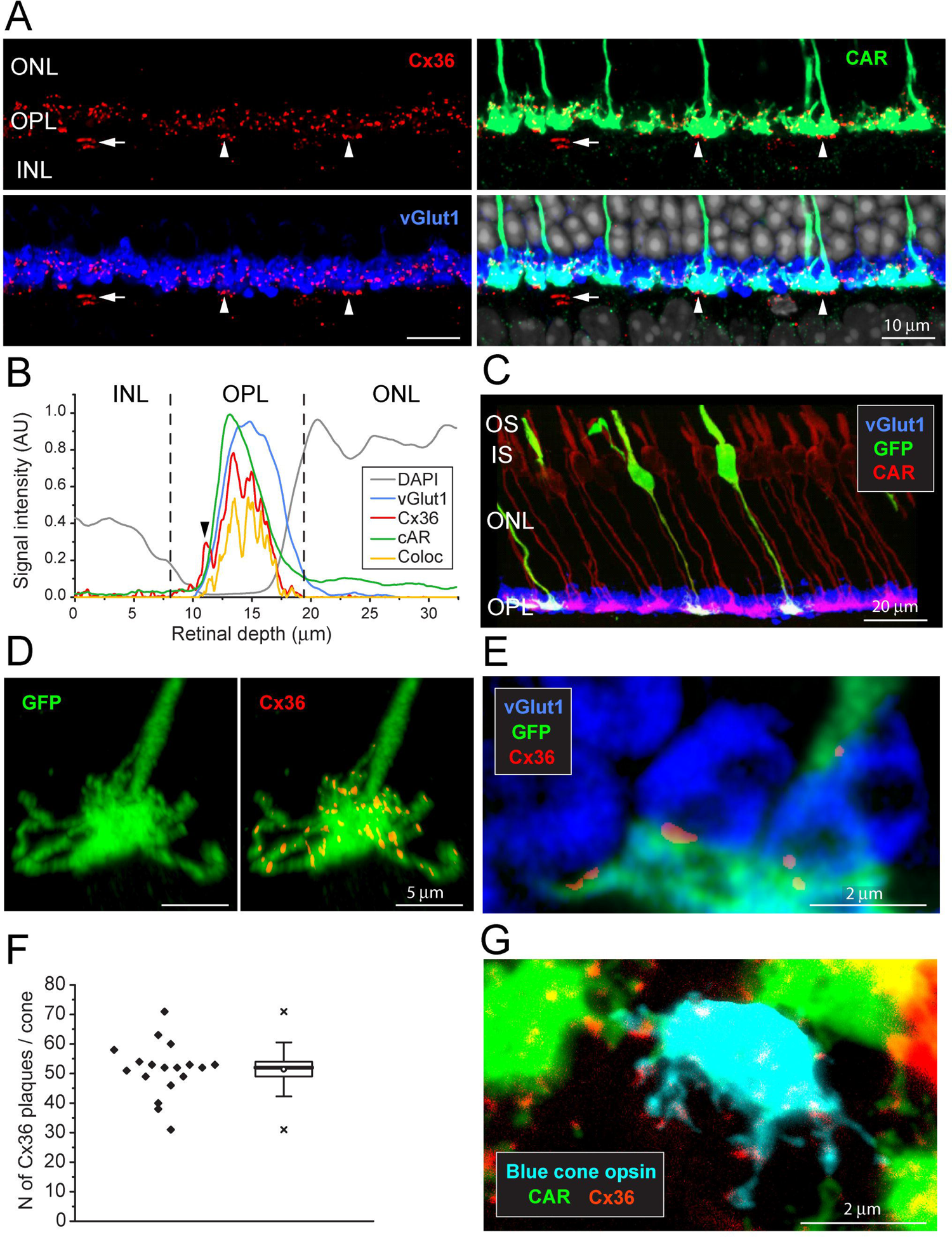
The Distribution of Cx36 in the OPL. **A.** Cx36 labeling in the OPL, confocal microscopy. Top left: numerous small Cx36 clusters (red) throughout the OPL, absent in the ONL. For all four panels: arrowheads, Cx36 under each cone pedicle, horizontal arrow, non-specifically labeled blood vessel. Top right: Cx36 clusters decorate the cone pedicles and their telodendria, labeled for cone arrestin (green). Lower left: Cx36 clusters are contained within the band of rod spherules, stained with an antibody against vGlut1 (blue). Lower right: 4 channels showing Cx36 (red), cone pedicles (green), and rod spherules (blue) all contained within the OPL. DAPI labeled nuclei (grey) show the well-organized ONL. Video 1 shows this data set. **B.** Analysis of A shows Cx36 was contained in the OPL with virtually no labeling in the ONL. Cx36 (red) is restricted to the band of cone pedicles (green) and rod spherules (blue). A small Cx36 peak outside the cone curve corresponds to the Cx36 under the cones (arrowhead). The non-specifically labeled blood vessels were removed for this analysis. Cx36 clusters colocalized with cone pedicles (orange) fit within the cone arrestin curve. **C.** EGFP labeled single cones (green) against a background of all cones stained for cone arrestin (red) and rod spherules (blue). **D.** 3D reconstruction of a single cone pedicle with many telodendria. Right, colocalized Cx36 clusters (red) on a single cone pedicle (green). Video 2 shows this data set. **E.** A single confocal section showing that rod spherules (blue) occur close to the Cx36 clusters (red) located on a cone pedicle (green). **F.** The number of Cx36 clusters, 51±8.9 (mean ± SD) on a single reconstructed cone pedicle, box shows quartiles, mean (circle), median (bold line), SD (whisker), min/max (x), n = 18. **G.** Blue cone pedicle (cyan), identified in a blue cone opsin Venus mouse line, had telodendria bearing Cx36 clusters (red), similar to those of green cones, stained for cone arrestin (green).

The cone telodendria rose above the level of the cone pedicles to contact the overlying rod spherules (Fig. 1A). In the OPL, there are many small Cx36 clusters associated with cone telodendria, often at their tips, relatively high (distal) in the OPL. Staining the rod spherules for a synaptic vesicle marker, such as the vesicular glutamate transporter, shows that Cx36 clusters are contained within the band of vGlut1 labeled rod spherules (Fig. 1A, Video 1). There was no Cx36 labeling in the outer nuclear layer (ONL). Cx36 clusters are also apparent underneath each cone pedicle (arrowheads, Fig. 1A), on processes previously identified as bipolar cell dendrites (Feigenspan et al., 2004; O’Brien et al., 2012; Raviola and Gilula, 1973).

We analyzed these data by plotting the intensity of Cx36 signal against depth in the retina (Fig. 1B). The vast majority of Cx36 clusters are contained within the OPL, within the bands of cone arrestin and vGlut1 labeling, which show the position of cone pedicles and rod spherules, respectively. On the left of the curve, there is a minor peak in the Cx36 channel (red line, black arrowhead), which falls outside the cone arrestin curve, corresponding to the Cx36 labeling of bipolar cell dendrites underneath each cone pedicle (Raviola and Gilula, 1973; O’Brien et al., 2012). Because this peak does not colocalize with cones, it was excluded from analysis and is not further considered in this paper. The area under the Cx36 curve indicates that an insignificant fraction (less than 0.2%) of Cx36 clusters are in the ONL. This is important because it rules out the presence of Cx36 gap junctions between rod somas and/or passing axons in the ONL, where they are packed together at high density.

### EGFP-labeled cones

When all the cones are labeled for cone arrestin, the complexity of the overlapping matrix makes it difficult to analyze individual cone pedicles with confidence. Therefore, we turned to a transgenic mouse line, *Opn4^cre^;Z/EG* (Ecker et al., 2010), where there is sparse labeling of a few individual cones (Fig. 1C). This made it possible to view individual EGFP-labeled cones against a background of all cones stained for cone arrestin. The immunofluorescence data were analyzed by extracting only the Cx36 clusters that colocalized with EGFP-labeled pedicles and reconstructing in 3D to find the number and location of Cx36 clusters on a single cone pedicle (Fig. 1D, Video 2). At each Cx36 cluster, there is an adjacent rod spherule (Fig. 1E), suggesting these structures are rod/cone gap junctions. There are approximately 51 ± 8.9 (mean, ± SD, n = 18) Cx36 clusters on each cone pedicle (Fig. 1F; Fig. 1 – source data 1), along the telodendria and over the upper surface of the cone pedicle.

### Blue cones – confocal microscopy

Blue cones initiate a specific color-coded pathway in the mammalian retina (Behrens et al., 2016; Dacey and Lee, 1994; Haverkamp et al., 2005; Kouyama and Marshak, 1992). Blue cones in the dorsal retina were stained in their entirety by use of a blue cone opsin Venus transgenic mouse line (Fig.1, suppl. 1). Alternatively, blue cones were identified using an antibody to blue cone opsin to stain the outer segment and then following the cone arrestin labelled axon down to the pedicle in a confocal series (Fig.1, suppl. 2, Video 3). In either case, a sparse mosaic of “true blue” cones were located in the dorsal retina (Nadal-Nicolás et al., 2020). We find that the pedicles of blue cones have telodendria bearing Cx36 labeled clusters in a manner indistinguishable from green cone pedicles (Fig. 1G; Fig.1, suppl. 2). Thus, it appears that blue cones also make numerous Cx36 gap junctions with rods.

### Cx36 clusters are located on rod spherules, close to the opening of the post-synaptic compartment

In high-resolution confocal images (63x objective, NA 1.4, Zeiss Airyscan), the Cx36 elements occur exclusively where cone telodendria contact the base of each rod spherule (Fig. 2A). Some rod/cone contacts have multiple Cx36 puncta (arrow, Fig. 2A). In these images, rod spherules appear as oval structures (blue) including two unlabeled compartments or holes, separated by the synaptic ribbon, immunolabeled for ribeye (white in Fig. 2B). The lower hole in the rod spherule of Fig. 2B contains the mGluR6-labeled (green) tips of rod bipolar cell dendrites and was thus identified as the post-synaptic compartment. The upper hole contains a single large mitochondrion, labeled for the mitochondrial translocase TOMM20 (red). To quantitatively determine the position of Cx36 labeled points, we captured vGlut1 labeled rod spherules and generated a mean structure by aligning, stacking and averaging the images from 18 complete rod spherules. The sites of individual Cx36 elements were marked and found to be located around the base of the mean rod spherule (Figs. 2C, 2D). A spline curve was fitted to the outline of the mean rod spherule and, after linearizing this curve, the density of Cx36 was plotted. The twin peaks of the resulting curve show that Cx36 (red) is distributed around the mouth of the synaptic opening, shown by a drop in the vGlut1 labeling (blue) (Fig. 2E). In the same region, the labeling for the cone signal (green) is high suggesting that Cx36 gap junctions occur at telodendrial contact points with rods. Two potential outliers, at approximately 3 o’clock and 9 o’clock (Fig. 2C), were actually located on other nearby rod spherules and so were excluded from the analysis. From the mean structure of 18 rod spherules, all Cx36 elements (2.5 ± 0.76/rod, (mean ± SD, n=45) are located at the base of the rod spherule, within 1 - 2μm of the opening to the post synaptic compartment.

**Figure 2.**
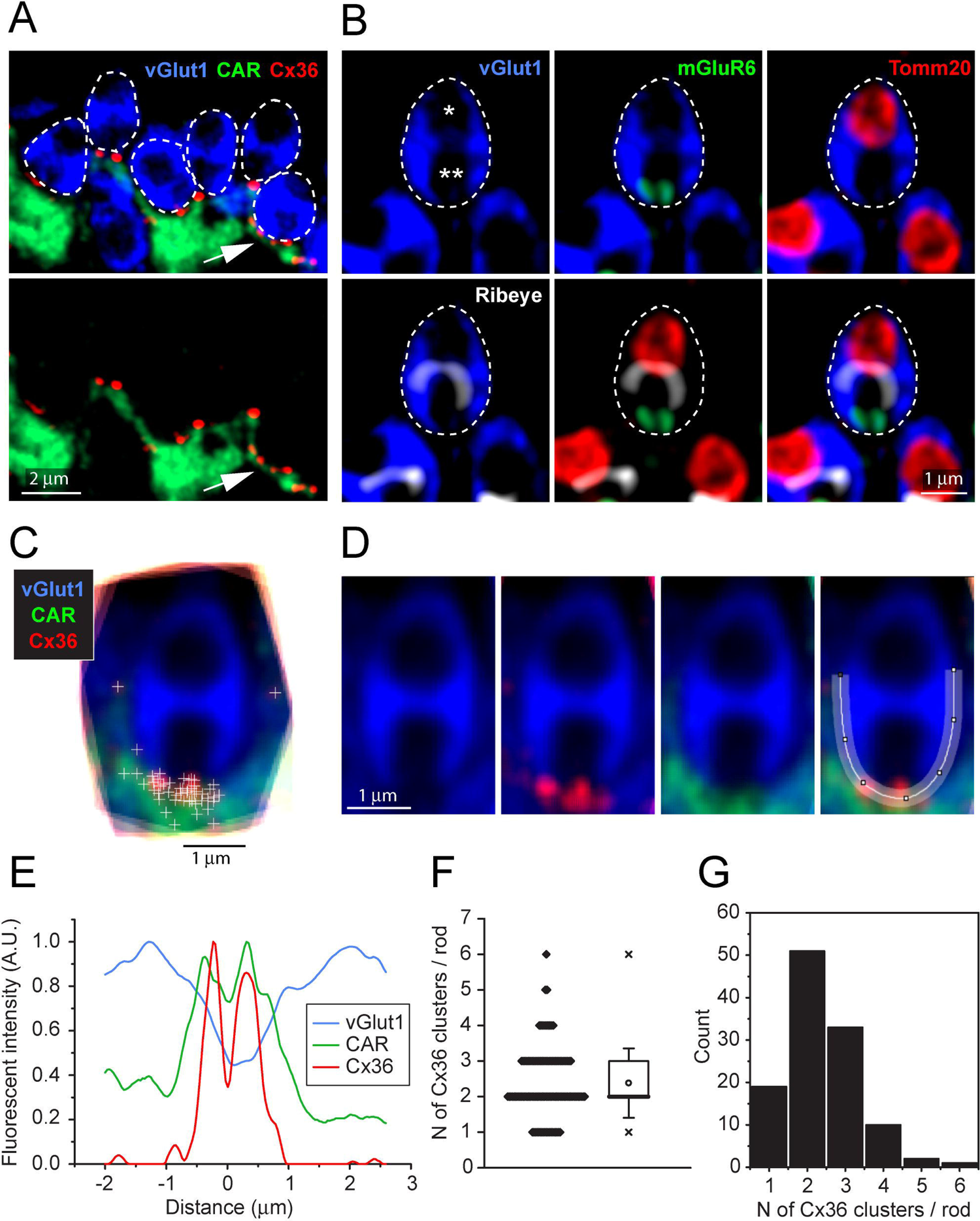
Cx36 clusters are found at the base of each Rod Spherule. **A.** Top, detail of OPL, rod spherules, stained for vGlut1 (blue) and outlined with dashed lines are located above and between cone pedicles labeled for cone arrestin (green). Cx36 clusters (red) are found at the base of rod spherules where they contact cones. Multiple Cx36 clusters occur at some contacts, arrow. Bottom, Cx36 clusters are located on cone telodendria. **B.** Top left, a rod spherule labeled for vGlut1 (blue), outlined with a dashed line, has two compartments (* and **). Top middle, bottom compartment is open at the base and contains two mGluR6 labeled rod bipolar dendrites, identifying the post-synaptic compartment. Top right, the upper compartment is filled with a large mitochondrion, labeled for TOMM20 (red). Lower left, the synaptic ribbon, labeled for ribeye (white), lies between the two compartments, arching over the post-synaptic compartment. Lower middle, the synaptic ribbon lies between the mitochondrion and the mGluR6 label. Bottom right, all four labels superimposed showing a typical rod spherule. **C.** 18 rod spherules, aligned and superimposed. Taking the rod spherule as a clock face, Cx36 clusters, (red with individual puncta marked +), are found at the base, 6 o’clock, along with cone telodendria (green). The two single +s at 3 o’clock and 9 o’clock are associated with other rod spherules and were excluded from the analysis. **D.** Average rod spherule showing Cx36 and cone telodendria at the base with a fitted spline curve. **E.** The distribution of label in each confocal channel along a linearized version of the spline curve. Where the vGlut1 label is low, indicating the opening to the post-synaptic compartment, the Cx36 (red) and cone (green) signals are high. The double peak for Cx36 indicates clusters on each side of the synaptic opening. **F.** Scatterplot showing the number of Cx36 clusters, presumed to be gap junctions, per rod spherule from a sample of 135 rods, mean 2.4 ± 0.97, box shows quartiles, mean (circle), median (bold line), SD (whisker), min/max (x). **G.** Data plotted as histogram showing the distribution of multiple Cx36 clusters per rod spherule, n=135.

To estimate the fraction of rods that are coupled to cones, we counted the number of Cx36 labeled points at the base of each rod spherule where there was contact with a cone, excluding any rod spherules that were not completely contained within the section. From three different samples, 116/116 rod spherules have cone contacts and Cx36 labeling close to the synaptic opening (Jin et al., 2020). Frequently, there are multiple Cx36 labeled points at the base of a single rod spherule; the number of Cx36 elements ranged from 1 - 6 with a mean of 2.4 ± 0.97 (mean ± SD, n=116 rods, total 276 Cx36 labeled points) (Fig. 2F, 2G; Fig. 2 – source data 1). Presuming that a Cx36 cluster indicates the presence of a gap junction, we conclude that every rod spherule was coupled to a nearby cone.

### Serial Blockface Electron Microscopy

The e2006 SBF-SEM dataset is derived from a block of mouse retina including 164 cone pedicles and thousands of rod spherules with a voxel size of 16.5 x 16.5 x 25 nm (Helmstaedter et al., 2013). Cone pedicles are easily recognized as the largest structures in the OPL and we were able to map them and register the resulting map with the data from Behrens et al (2016) (Fig. 3A). From these data, we could locate the blue cone pedicles previously identified by their selective contacts with blue cone bipolar cells (Behrens et al., 2016). Rod spherules were also easily identified as the numerous round and compact structures with prominent post-synaptic inclusions in the synaptic invagination. In this dataset, rod spherules are 2-3μ they are massed above and surrounding the cone pedicles (Fig. 3B), usually oriented with the synaptic invagination at the base.

**Figure 3.**
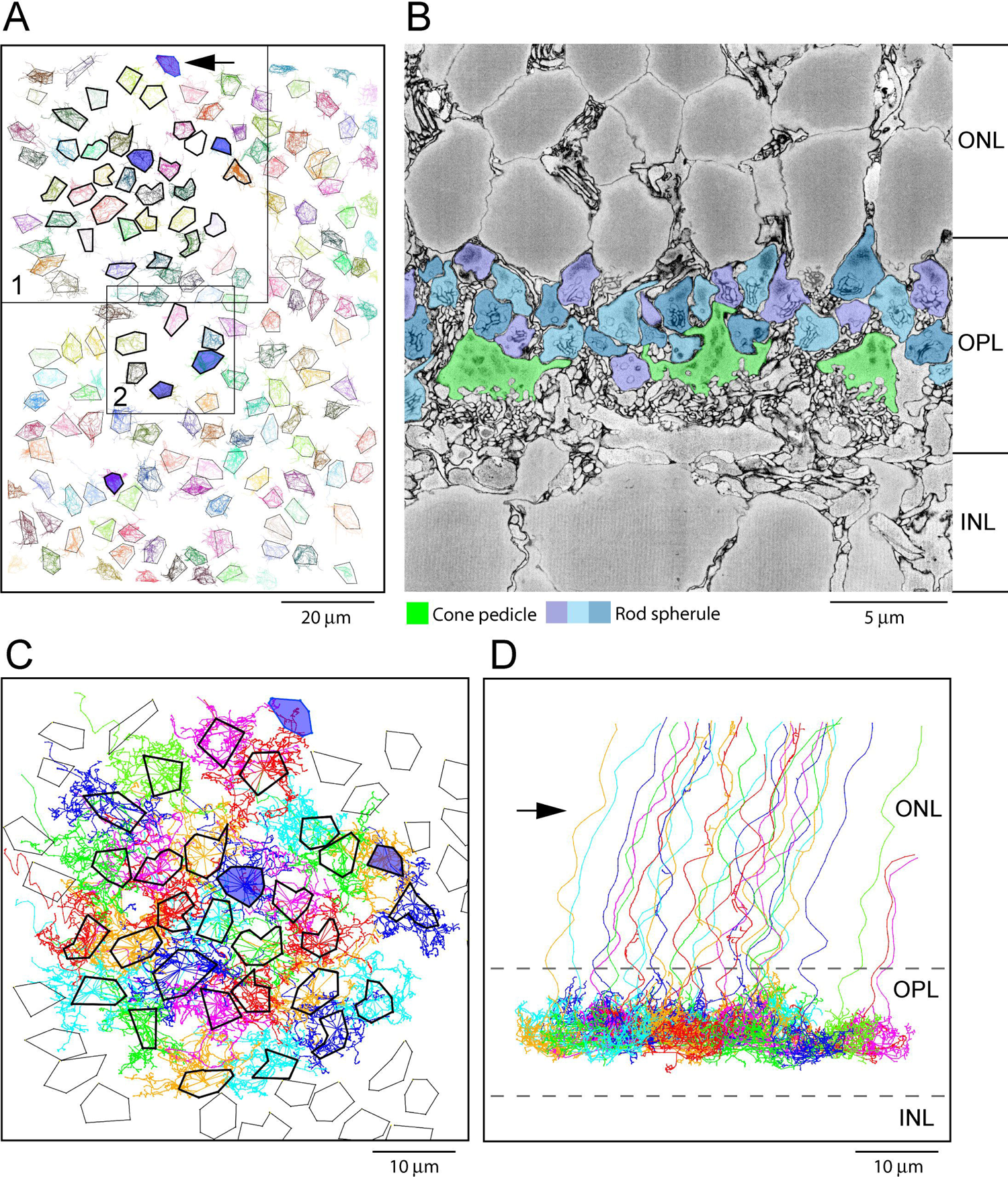
Map of Cone Pedicles shows Telodendria Overlap. **A.** Map of 164 cone pedicles in e2006, boxes 1 and 2 (low density, also Fig. 4, suppl. 1) show reconstructed areas, dark outlines show reconstructed cone pedicles, 6 blue cone pedicles filled dark blue (Behrens et al., 2016). **B.** Vertical section from e2006 showing rod spherules (3 shades of blue), above and in between cone pedicles (green). **C.** Skeletons of all 29 reconstructed cone pedicles (thick outlines) showing overlapping telodendrial fields, individually colored. Thick outlines show solid part of cone pedicles. **D.** Projection of C showing cone pedicles restricted to the OPL and axons ascending through the ONL.

### Skeletons show Many Contacts between Cone Telodendria and Rod Spherules

To assess potential gap junctional contacts between cone pedicles and rod spherules, we chose a patch of 29 adjacent cones to skeletonize, meaning we followed their processes to nearby contacts or termination. This patch included 13 central cone pedicles surrounded by a ring of 16 additional cone pedicles (Fig.3A). Some calculations were based on the central 13 cones to avoid edge artefacts. All skeleton data are provided in Fig. 3 – source data 1 and summarized in Table 1, Appendix 2. This area contained one blue cone pedicle but, in addition, we analyzed all the blue cone pedicles in the data set, as identified by Behrens at al (2016) (summarized in Table 2, Appendix 2). We also examined several other locations, including areas with a sparse distribution of cone pedicles, to be sure we did not select an atypical area (Fig. 3A, Fig. 4, suppl. 1, C & D). The mosaic of cone pedicles with their overlapping telodendria is plotted in Fig. 3C. Cone axons were followed into the ONL and a projection shows that the telodendria are contained within the upper part of the OPL (Fig 3D).

**Figure 4.**
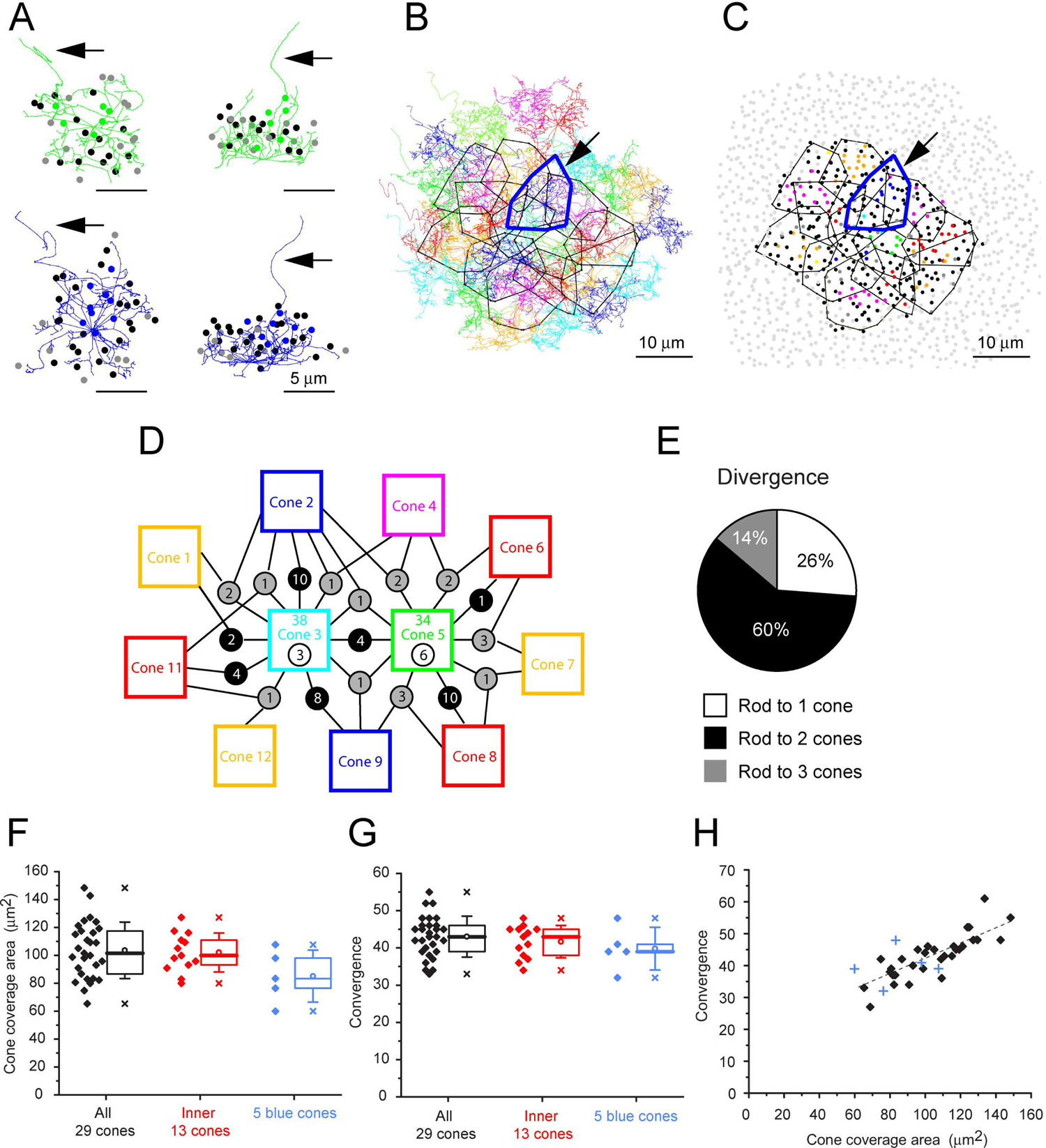
Cone Skeleton Analysis shows Cones contact all nearby Rod Spherules. **A.** Skeletons of one green cone (cone 5, green) and one blue cone (cone 2, blue) in wholemount view and projected. Arrows show ascending axon. The position of each contacted rod spherule is marked by a dot, color coded the same color (green or blue) if the contacts are exclusive to this cone pedicle, black, contacts 2 cones, grey, contacts 3 cones. **B.** Telodendrial fields of 29 reconstructed cone pedicles, each color-coded, central 13 outlined by polygons, arrow points to blue cone (cone 2). **C.** Outlines of central 13 cone pedicles showing all rod spherule contacts, color coded, exclusive to one cone, black, contacts with 2 cones, dark grey, contacts with three cones. Light grey, rod spherules outside the range of the central 13 cone pedicles. **D.** Simplified skeleton map showing contacts of two cone green pedicles (cones 3 and 5), the box contains the cone identity, the total number of rod contacts (above) and the exclusive number of rods which contact this cone only. Circles indicate the number of rod spherules shared between cones linked by lines to neighbors. Cone 2 is a blue cone, which shares rod contacts with neighboring cones. **E.** Pie chart showing distribution of cone contacts per rod spherule. **F.** Convergence, rod contacts per cone for all 29 reconstructed cone pedicles, inner 13 and 5 blue cones. Box shows quartiles, mean (circle), median (bold line), SD (whisker), min/max (x). **G.** Cone pedicle telodendrial field area for all 29 reconstructed cone pedicles, inner 13 and 5 blue cones. Box shows quartiles, mean (circle), median (bold line), SD (whisker), min/max (x). **H.** Convergence vs cone pedicle area. Line of linear regression shows some tendency for larger cone pedicles to contact more rods. +, blue cones

**Table 1.**

|  | Whole cones | Inside cones | p-value |
| --- | --- | --- | --- |
| N of cones | 29 | 13 |  |
| N of rods | 811 | 361 |  |
| Rods / cone | 28.0 | 27.8 |  |
| Convergence |  |  |  |
| mean ± s.d. | 43.0 ± 5.4 | 41.7 ± 4.2 | 0.40 |
| median (Q1-Q3) | 43 (39 - 46) | 43 (38 - 45) |  |
| Divergence | 1.54 ± 0.63 | 1.89 ± 0.64 | 2.5 x 10<sup>-17</sup> |
| Conv. / Div. | 28.0 | 22.1 |  |
| Cone coverage area* |  |  |  |
| mean ± s.d. | 103 ± 20 | 102 ± 14 | |
| median (Q1-Q3) | 100 (86.7 - 117) | 99.9 (93.0 - 111) |  |
| Area covered by cones* | 1920 | 890 |  |
| Cone density** | 0.0151 | 0.0146 |  |
| Coverage | 1.56 | 1.49 |  |
| N of shared rods |  |  |  |
| mean ± s.d. | 6.23 ± 4.67 | 5.85 ± 4.42 | |
| median (Q1-Q3) | 4 (2 - 8) (N=79) | 5 (2 - 9) (N=61) | 0.63 |

Unit is $\mu\text{m}^2$ and $\mu\text{m}^2$ .

**Table 2.**

|  | Blue cones | Green cones | p-value |
| --- | --- | --- | --- |
| N of cones | 5 | 31 |  |
| Convergence |  |  |  |
| mean ± s.d. | 39.8 ± 5.11 | 43.0 ± 6.9 | 0.31 |
| median (Q1-Q3) | 39 (39 - 41) | 43 (39 - 46) |  |
| Cone coverage area* |  |  |  |
| mean ± s.d. | 85.1 ± 18.6 | 104 ± 22 | |
| median (Q1-Q3) | 83.4 (76.4 - 98.0) | 105 (84.8 - 120) | 0.081 |
| N of shared rods |  |  |  |
| mean ± s.d. | 6.56 ± 6.34 | 6.26 ± 4.14 | |
| median (Q1-Q3) | 6 (3 - 9) (N=16) | 3.5 (2 - 10.8) (N=69) | 0.86 |

\*Unit is $\mu\text{m}^2$ .

We started our analysis with a single cone pedicle (cone 5), near the center, which had a complex but typical field of telodendria approximately 10µm in diameter, and we marked the position of every rod spherule in contact with this cone (Fig. 4A). Cone 5 has contacts with 34 rod spherules which included every rod spherule within the telodendrial field. In a projection, it can be seen that the telodendria extend laterally and above the cone pedicle, reaching up to the overlying rod spherules (Fig. 4A). Many, but not all, telodendria terminate at the base of a rod spherule. The skeleton of a blue cone pedicle (cone 2) is similar (Figs. 4A, bottom and 4B, arrow).

Outlining the telodendrial fields and adding the rest of the cone pedicles shows a dense matrix of overlapping telodendria (Fig. 4B) each with an area of 103 ± 20 µm (mean ± SD, n=29, Fig. 4F; Fig. 4 – source data 1) and a coverage estimated at 1.56 (n=29 cones). Each cone contacts 43 ± 5.4 rod spherules (mean ± SD, n=29 cones) (Fig. 4C & G; Fig. 4 – source data 1), including every rod spherule within its field, typically at the base. Plotting convergence (number of rod spherule contacts per cone) against the pedicle area shows a slight trend for the largest cone pedicles to contact more rods spherules (Fig. 4H; Fig. 4 – source data 1).

Examining these contacts from the perspective of the rod spherules, we coded the overlying rod spherules in the same color as each cone pedicle. Rod spherules with contacts from two or three of the central 13 cone pedicles are marked black or dark grey respectively with non-contacted rod spherules marked as light grey. Viewing the resulting map quickly makes the point that most rods are contacted by two or three cone pedicles (Figs. 4C; Fig. 4, suppl. 1A). Omitting the cone pedicles (except for one example) and all rod spherules in contact with those cones, leaves only an annulus of rod spherules outside the field of reconstructed cones and demonstrates the almost total absence of rods with no cone contacts (Fig. 4, suppl. 1, B). We found 3/811 (0.4%) rods without cone contacts within the field of 29 cones, an insignificant fraction. As a precaution, we also checked a sparse patch with a rare hole in the cone mosaic (Fig 4. Suppl. 1, C and D). Even here, more than 95% of rods received cone contacts.

If each cone contacts every rod spherule within its telodendrial field, then in areas where the cone telodendria overlap, each rod must receive contacts from multiple cones. This is clearly the case, and, in fact, most rods receive contacts from two, three or, rarely, four cones. A simplified map for two adjacent cones is shown in Fig. 4D, while the contact map for all 29 cones is shown in Fig. 4 supplement 2A. Of approximately 40 rods in contact with each cone pedicle, only a few have exclusive contacts with a single cone (7.2 ± 3.4, mean ± SD, n=13 central cone pedicles). Most rods receive contacts from several cone pedicles while adjacent cone pedicles share as many as 23 rod spherules, (mean ± SD = 6.2±4.7, n=79 cone pairs) (Fig. 4 suppl. 2B; Fig. 4 – source data 2). There is a pronounced edge effect because the outermost ring of cone pedicles lacks the overlap from further unanalyzed cones: to avoid this, we analyzed the rod contacts of the central 13 cone pedicles. In this area, rods with multiple cone contacts were the norm; 74% of rod spherules received contacts from two or three cones, yielding a rod to cone divergence of 1.9. (Fig. 4E). This reflects the density of the telodendrial network and the large number of rod/cone gap junctions in the OPL. Clearly, each rod spherule has the potential to make several contacts with close-by cones, in agreement with the confocal data showing the distribution of Cx36.

### Blue Cone Skeletons

The skeleton of a blue cone, defined by its bipolar cell contacts (Behrens et al., 2016; Nadal-Nicolás et al., 2020), is also shown in figure 4A and the analysis of all five blue cones from the dataset of Behrens et al, (2016) is presented for comparison with green cones (Fig. 4F & G). The blue cone data are summarized in Table 2, Appendix 2. The telodendrial area of blue cones may be slightly smaller, 85 ± 19μm^2^ (mean ± SD, n=5) (Fig. 4F) but the difference is not significant due to the small sample size. By all other measures, blue cones are indistinguishable from green cones. Blue cones contact all rod spherules, (40 ± 5.1, mean ± SD, n=5) within their telodendrial field, most of which also receive contact from adjacent green cones. Thus, there is no evidence for color selective rod contacts.

### Segmentation and Reconstruction

We segmented several cone pedicles and the rod spherules contacted by each cone pedicle. The contact sites, where the cell membrane of a cone telodendron merged with the cell membrane of a rod spherule, were highlighted. These contact pads could then be superimposed on either the cone pedicle or the surface of the overlying rod spherules. Typically, the telodendrial contacts, often from more than one cone, appear as arcs, close to the synaptic opening of rod spherules (Fig 5, A-D) (Kolb, 1977; Smith et al., 1986; Tsukamoto et al., 2001). Confocal analysis also demonstrated multiple cone contacts with a single rod spherule (Fig. 5E).

**Figure 5.**
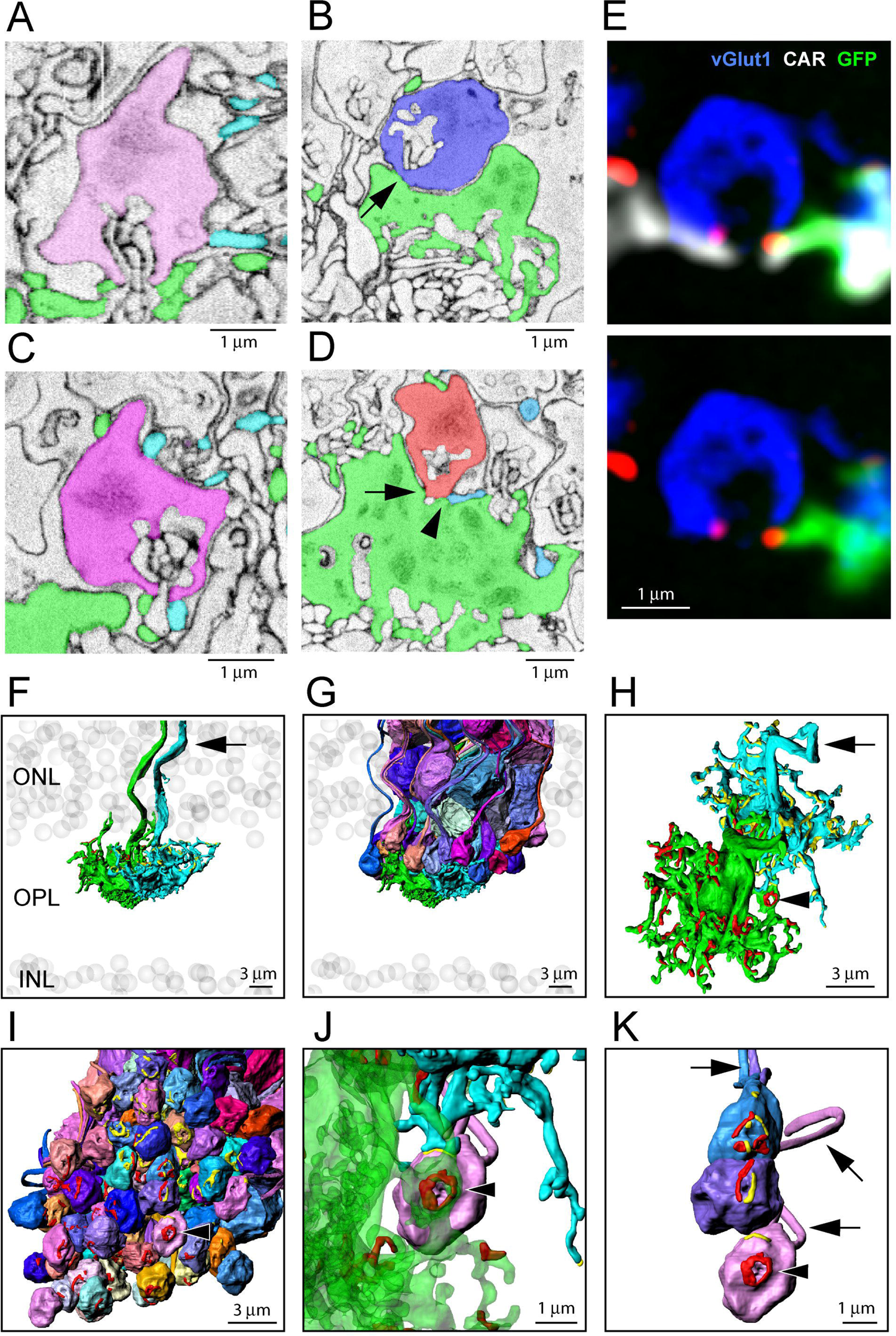
Segmentation and 3D Reconstruction e2006: Cone Telodendria contact Rod Spherules close to the Mouth of the Synaptic Opening. **A.** Single SBF-SEM section, telodendria from 2 cones (green or cyan), rod spherule (pink), in the plane of the synaptic opening with cone contacts on both sides. **B.** Single SBF-SEM section, rod spherule (blue), post-synaptic inclusions show it is close to the synaptic opening, 25 nm x 4 sections = 100 nm away. The rod spherule sits on the roof of a cone pedicle (green) with a contact, shown by the arrow, where membranes merge. **C.** Single SBF-SEM section, rod spherule (magenta) with contacts at the synaptic opening from 2 different cones (green or cyan). **D.** Single SBF-SEM section, rod spherule (orange), contacts (arrow) the roof of a cone pedicle (green) with a telodendrial contact (arrowhead) from a second cone (cyan). Post-synaptic inclusions in the rod spherule indicate the contacts are close to the synaptic opening, 25 nm x 9 sections = 225 nm away. **E.** Top, confocal microscopy, single optical section, shows Cx36 clusters (red) where telodendria from 2 different cones (white for cone arrestin, labels both cones; green for a single EGFP labeled cone) contact rod spherule (blue), at the mouth of the synaptic opening. Bottom, cone arrestin signal turned off for clarity, showing only one cone is labeled for EGFP. **F.** 3D reconstruction from e2006, 2 adjacent green cone pedicles (cone 5, green, and cone 3, cyan), ghost cell bodies (grey) show limits of ONL and INL, arrow marks ascending cone axons. **G.** Same 2 cone pedicles with all 66 (= 38 + 34 – 6 shared) reconstructed rod spherules contacted by these 2 cones. **H.** Rotated view, looking down at top surface of same 2 cone pedicles, contact pads with rod spherules marked in red or yellow for each cone. **I.** Rotated view, looking up at the bottom surface of all rod spherules in contact with these 2 cones, contact pads in red or yellow, arrow head marks a single rod spherule magnified in J and K. **J.** Single rod spherule (pink), green cone pedicle (cone 5), with adjusted transparency, with contact pad (red) encircling the synaptic opening at the base of the rod spherule (arrowhead). The second cone (cone 3, cyan) also contacts this rod spherule nearby (yellow), approximately μm from the synaptic mouth. **K.** Detail, showing the bottom surface of 3 adjacent rod spherules which receive contacts close to the synaptic opening from both cone pedicles, contact pads in red or yellow, arrows show rod axons, arrowhead indicates same rod spherule as J. Video 5 shows this data set.

The reconstruction of adjacent cones 3 & 5 shows a complex field of telodendria extending laterally and upwards (distally) from the pedicles (Fig. 5F). Their telodendria interdigitate, mostly avoiding each other; the sparse cone-to-cone contacts will set a limit on cone/cone coupling. We also reconstructed all the rod spherules in contact with these cones (Fig. 5G, Videos 4 & 5). Most of the overlying rod spherules have telodendrial contacts (Figs 5A & C) but a few make direct contacts with the upper surface or roof of the cone pedicle (Fig. 5B & D, Fig. 5 suppl. 1A). When the contact sites with rods are displayed on the cone pedicles, the appearance is similar to the 3D reconstruction of confocal material showing the distribution of Cx36 clusters on a single cone pedicle (Figs. 1D & 5H). When the contact sites are displayed on the rod spherule lower surfaces, it is obvious that most telodendrial contacts are very close to the mouth of the synaptic invagination, often forming a curved line or horseshoe around it (Fig. 5I, J), within 1-2µm. Neighboring cone pedicles often contact the same rod spherules. At such sites, the contact pads from both cone pedicles can be found, often interspersed around the synaptic mouths of the rod spherule (Fig. 5J & K, Video 4 & 5). The cone contact sites are consistent with the location of Cx36 near the mouth of the synaptic invagination and thus indicate the potential presence of rod/cone gap junctions (Fig. 5E). However, a contact pad may exceed the extent of a gap junction and therefore does not predict its size (see below).

Of the cone pedicles that were completely reconstructed, cones 5, 3 and 2 (a blue cone), had 57, 59 and 54 contact pads respectively. Thus, the number of contact pads is close to the number of Cx36 clusters per cone pedicle, (51 ± 8.9, mean ± SD), from the confocal analysis, and this suggests that most of the contact pads include gap junction sites. There were a few examples where a cone telodendron contacts a rod spherule far from the synaptic invagination, often in transit to another location, and these can show large areas of contact. But these sites are not associated with Cx36 labeling in the confocal data. In other words, these are incidental contacts of passage, not gap junctions, and they were excluded from our analysis.

There are a few anomalies in the OPL: some rod spherules sit directly astride a cone pedicle and receive few or no telodendrial contacts. Instead, they make direct contacts with the roof of the cone pedicle (Fig. 5 suppl. 1). In addition, the lowest (most proximal) row of rods in the ONL do not have axons or spherules (Li et al., 2016). Instead, the synaptic machinery is included in the lowest crescent of the cell body, adjacent to the OPL. These low rods represent the most distal synaptic structures in the OPL, yet they still receive contacts from cone telodendria (Fig. 5 suppl. 2). Finally, there are a few rod spherules below (proximal to) the level of the cone pedicles. These are often inverted so the mouth is on top and this is the site of telodendrial contacts (Fig. 5 suppl. 3). Thus, in every case, including these anatomical variations, cone contacts occur close to the rod synaptic opening, coincident with the location of Cx36. These sites very probably contain Cx36 rod/cone gap junctions.

### Blue Cone Segmentation

Based on the work of Behrens et al. (2016), who identified the cones in contact with the blue cone bipolar cell (b9), we could locate the blue cones in the map of cone pedicles. The telodendrial field of cone 2, a blue cone, contacts 41 rod spherules and overlaps substantially with the surrounding cones (Fig. 5, suppl.4A). This blue cone pedicle made the same types of rod contacts as other cones, including telodendrial contacts, roof contacts and contacts with the rods in the bottom row of the ONL (Fig. 5, suppl.4B). Reconstructing in 3D shows typical contact pads around the mouth of the overlying rod spherules with alternating contributions from multiple cones, including this blue cone. (Fig. 5, suppl.5). Thus, blue cone pedicles also make rod/cone gap junctions and both blue and green cones can make gap junctions with the same rod. We found no evidence of color-specific coupling.

### SBF-SEM, Singer dataset (eel001)

While the SBF-SEM data analyzed above provided extensive information about close appositions between cone pedicles and rod spherules, the sample preparation was not appropriate to resolve definitive gap junctions in the tissue. To demonstrate that the contact pads described above represent the presence of gap junctions, we also examined rod/cone contacts in a SBF-SEM dataset (eel001) in which synapses and rod/cone gap junctions were visible. In Fig. 6 A & B, a roof contact between a cone pedicle and an overlying rod spherule is shown. There is clear separation between the cell membranes until a darkly stained chromophilic area where the membranes merge indicating a gap junction approximately 350nm in length. The post-synaptic inclusions of the rod spherule show that this site is close to the opening of the post-synaptic compartment. We have shown above that Cx36 occurs at such contacts between rods and cones and therefore conclude that this demonstrates the presence of a gap junction.

**Figure 6.**
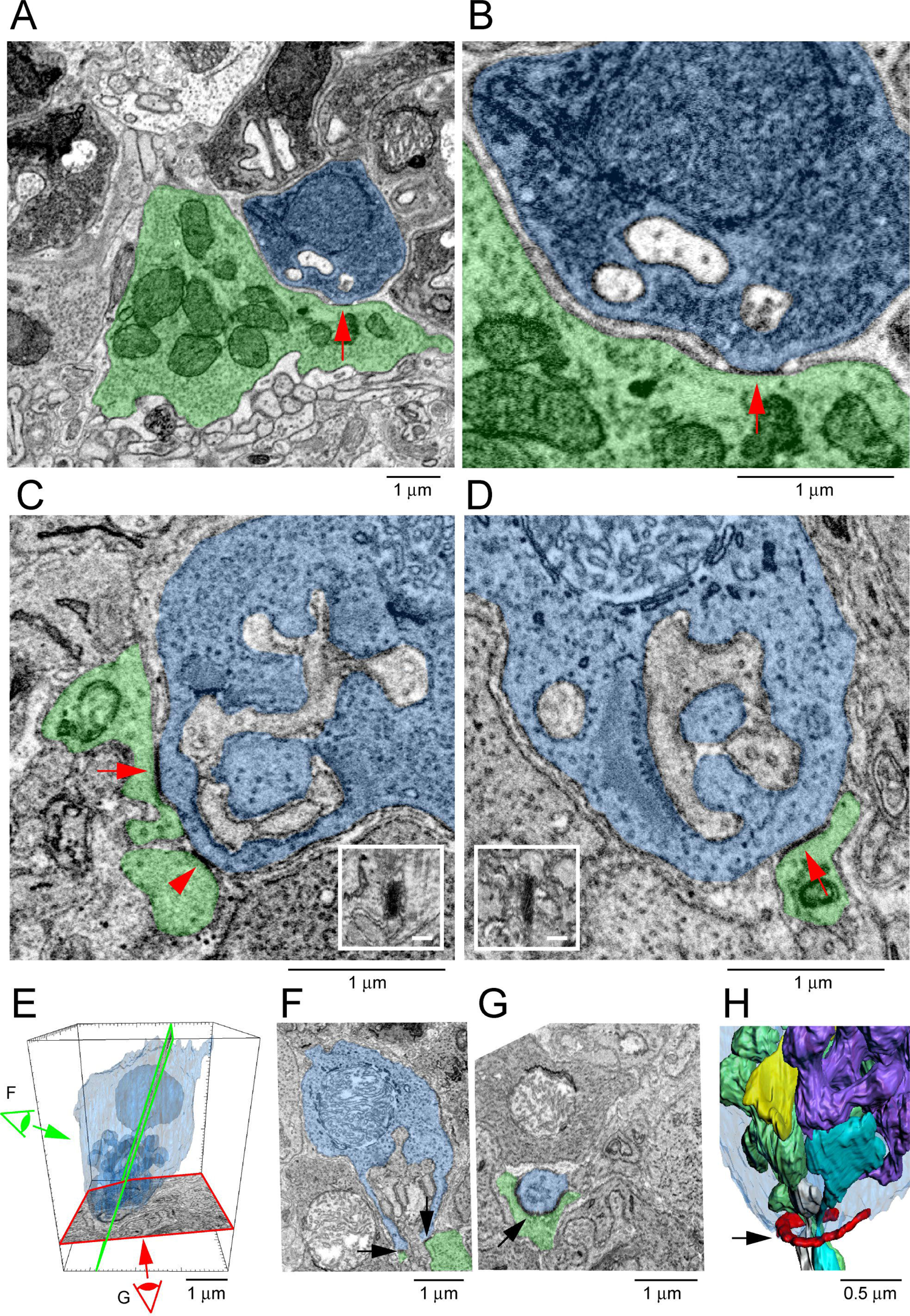
Segmentation and 3D reconstruction shows Gap Junctions at Rod/Cone Contacts. **A.** Singer dataset, SBF-SEM, a rod spherule (blue) nestled on the roof of a cone pedicle (green) with a densely stained gap junction contact (arrow). **B.** Singer dataset, SBF-SEM, detail showing a gap junction (arrow) with merged membranes Note, there is clear separation except at this point. Post-synaptic inclusions indicate the gap junction is close to the synaptic opening of the rod spherule. **C.** FIB-SEM: Dense gap junction staining at contact points (arrows) between cone telodendria (green) and a rod spherule (blue). Inset: rotated en face view of one gap junction to measure length. **B. D.** FIB-SEM: second example (arrow) of a dense gap junction at a cone contact (green) with a rod spherule (blue). Inset: en face view of the gap junction. Note green cone telodendrial contact exceeds the gap junction staining. **C. E.** FIB-SEM: Overview of a rod spherule (blue) with a transparent reconstruction over the image planes in F and G. Note the large mitochondrion towards the top of the rod spherule. **D. F.** Image plane of the rod spherule (blue) through the synaptic opening with a small gap junction on each side (arrows) at a cone contact (green). **E. G.** Image plane close to the base of the rod spherule (blue) reveals the 2 small gap junctions in F are part of a single large gap junction. There is a horseshoe-shaped gap junction (arrow) close to the synaptic opening where a cone telodendron (green) wraps around the base of the rod spherule. Large Cx36 horse-shoe shaped Cx36 clusters like this are easily observed by confocal microscopy. **F. H.** FIB-SEM 3D reconstruction of the rod spherule (transparent blue) showing a single large gap junction (red, arrow) around the synaptic opening at the base. Post-synaptic inclusions are also rendered with 2 rod bipolar dendrites (grey and cyan) and 2 horizontal cell processes (green and purple). The synaptic ribbon is shown in yellow, partially obscured by the post-synaptic processes. Video 6 shows this data set.

### FIB-SEM to Visualize Gap Junctions

We also obtained two FIB-SEM data sets of mouse OPL (FIB-SEM 1 and FIB-SEM 2) with an isotropic resolution of 4nm, permitting the direct visualization of gap junctions. Contacts from cone telodendria at the base of each rod spherule show darkly stained gap junctions consistent with the location of Cx36 labeling (Fig. 6C, D). It should be noted that the area of telodendrial contact is frequently larger than the size of the gap junction (Fig. 6D). Thus, while contact pads in the SBF-SEM e2006 images indicate the potential for a gap junction (Fig. 5 H-K), they do not predict its size.

Sometimes, what appear as small separate gap junctions in a single section merge in nearby sections to form one large continuous gap junction. An example is shown in Fig. 6E-H; two small gap junctions on either side of the synaptic opening are part of a horseshoe structure similar in shape to many of the reconstructed contact pads. Reconstructing in 3D, shows one large gap junction encircling the base of the rod spherule with a length of approximately 1.5µm (Fig. 6H, Video 6). This curved structure is highly reminiscent of the curved gap junction strings revealed by freeze-fracture EM (Raviola and Gilula, 1973). In the OPL, a few large gap junctions of a similar shape, >1µm long, were readily apparent in the confocal view of wholemount retina labeled for Cx36 (Jin et al., 2020).

We were able to rotate these gap junction contacts in 3D and estimate the gap junction size from the en face view, as well as the distance from the mouth of the post-synaptic compartment. We analyzed 42 complete rod spherules with a total of 135 rod/cone gap junctions. These data are summarized in Table 3, Appendix 2. In the two FIB-SEM datasets, 42/42 rod spherules have gap junctions close to the mouth of the synaptic compartment consistent with the location of Cx36 (Figs 2A-E). We could not always trace the contact back due to the small volume of these high-resolution data sets, but when it was possible, in 112 cases, 100% of the gap junction contacts were identified as a cone. The number of gap junctions per rod spherule ranged from 1-6 with a mean of 3.2±1.2, (mean, ± SD, n=135, Fig. 7A; Fig. 7 – source data 1). The mean distance from the synaptic mouth was 0.69 ± 0.64µm, (mean, ± SD, n=135), but this is a skewed distribution. 84% of gap junctions lay within 1µm of the synaptic mouth with a median distance of 0.44µm; 94% were within 2µm (Fig. 7B & C; Fig. 7 – source data 1), coincident with the position of cone contacts and the location of Cx36 labeling (Fig 2C-E). There were 8 gap junctions, 6% of the total, which fell outside this range. When these outliers were traced back, the contacts were identified as cones in every case, so despite their distant location relative to the rod synaptic mouth, they were identified as rod/cone gap junctions.

**Figure 7.**
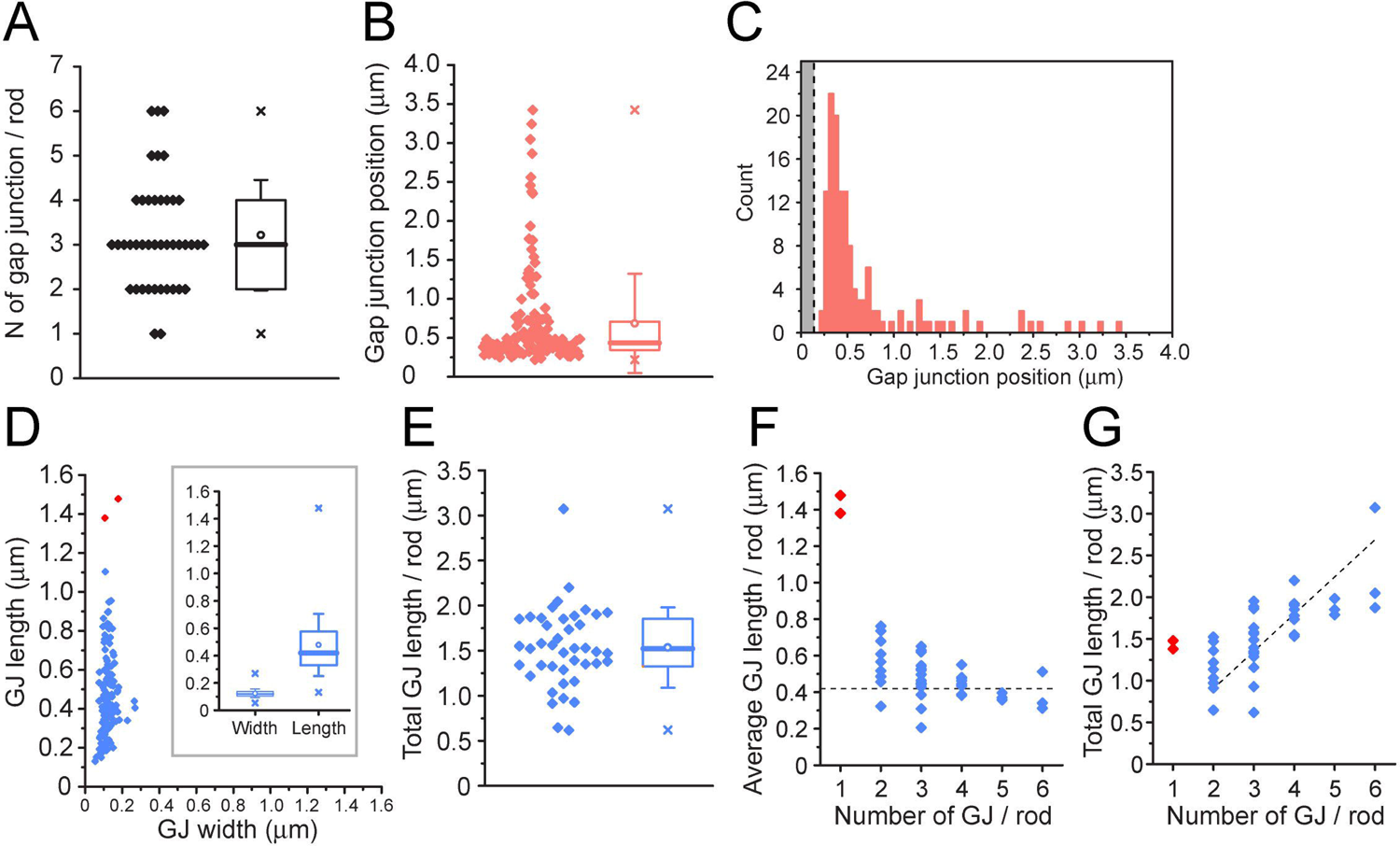
Quantitative Analysis of 42 Reconstructed Rod Spherules from FIB-SEM. **A.** The number of gap junctions per rod spherule ranged from 1-6 (3.2 ± 1.2, mean ± SD, n=42). Box shows quartiles, mean (circle), median (bold line), SD (whisker), min/max (x). **A. B.** Gap junction distance from the rod spherule synaptic opening, skewed distribution, median, 0.44 μm. Box shows quartiles, mean (circle), median (bold line), SD (whisker), min/max (x). **A. C.** Histogram plots of gap junction position, grey area shows diameter of the synaptic opening, (0.28±0.024μm, n=42 rod spherules) **B. D.** Gap junction length vs width for 135 gap junctions from 42 rod spherules. Note restricted width compared to much greater variability in length, consistent with string-like structure. Box shows quartiles, mean (circle), median (bold line), SD (whisker), min/max (x) for width and length. Outliers in red are from rod spherules with a single large gap junction. **C. E.** Total gap junction length per rod for 42 rod spherules. Box shows quartiles, mean (circle), median (bold line), SD (whisker), min/max (x). **D. F.** Average gap junction length plotted as a function of the number of gap junctions per rod shows that if there is only one gap junction per rod spherule, it is a large one. The 2 longest gap junctions from D were both singles (red, same two as in D). These are easily observed when labeled for Cx36 using confocal microscopy (Fig. 1, suppl 2; Jin et al., 2020). The dashed line is the median gap junction length (0.42μm) and it runs through the data for multiple gap junctions per rod. In other words, each additional gap junction is approximately a standard size. **A. G.** Total gap junction length per rod spherule as a function of the number of gap junctions per rod shows that the single gap junctions (red) are outliers with a large area. Total gap junction area tends to rise with the number of gap junctions. The straight line of linear regression shows the effect of adding a standard mean gap junction length (0.44μm) each time and runs through all the data except for the singles (red). Fitted line is y = 0.44x; (R^2^ = 0.33).

**Table 3.** FIB-SEM analysis

|  |  |
| --- | --- |
| N of rods | 42 |
| N of GJ / rod |  |
| mean ± s.d. | 3.21 ± 1.23 |
| median (Q1-Q3) | 3 (2 - 4) |
| GJ position* |  |
| mean ± s.d. | 0.686 ± 0.635 |
| median (Q1-Q3) | 0.435 (0.345 - 0.703) |
| Synaptic mouth radius* | 0.138 ± 0.121 |
| GJ width* |  |
| mean ± s.d. | 0.123 ± 0.032 |
| median (Q1-Q3) | 0.120 (0.103 - 0.138) |
| GJ length* |  |
| mean ± s.d. | 0.477 ± 0.227 |
| median (Q1-Q3) | 0.419 (0.332 - 0.568) |
| Total GJ length / rod* |  |
| mean ± s.d. | 1.53 ± 0.44 |
| median (Q1-Q3) | 1.52 (1.33 - 1.85) |

Unit is \*( $\mu\text{m}$ ) and \*\*( $\mu\text{m}^2$ ).

The identity of the dense chromophilic material visible in EM pictures of gap junctions is unknown but it may consist of gap junction proteins (connexins) in addition to scaffolding proteins and modulatory subunits that are thought to make up a gap junction complex (Lasseigne et al., 2021; Nagy et al., 2018). Freeze-fracture EM analysis of retinal gap junctions has shown that they exist in several distinct forms, from plaques with a crystalline array to sparse linear forms such as ribbons and strings (Kamasawa et al., 2006). En face, the rod/cone gap junctions had a relatively uniform width (∼120nm), suggesting a standardized component with a variable length such as a string, as previously reported in freeze fracture studies of the OPL (Raviola and Gilula, 1973). This is also consistent with the low fluorescent intensity of Cx36 in the OPL compared to the IPL, where gap junctions occur predominantly in the form of bright Cx36 clusters representing dense two-dimensional arrays of connexons (Kamasawa et al, 2006).

The mean length of a rod/cone gap junction was 480nm ± 230nm, (mean ± SD, n= 135, 42 rods, Fig. 7D; Fig. 7 – source data 1). Raviola and Gilula (1973) reported the photoreceptor gap junctions as strings of single particles with a spacing of 10nm. Therefore, the average rod/cone gap junction contains a string of 48 connexons. The total gap junction length per rod spherule was 1.5 µm ± 0.44 µm (mean ± SD, n=42, Fig. 7E; Fig. 7 – source data 1) or 150 channels. The largest individual gap junctions occurred when there was only one gap junction per rod spherule (Fig.7, D, F & G, red points; Fig. 7 – source data 1); these were outliers with a length close to the mean total length and they were visible in confocal images as large curved Cx36 labelled structures (Jin et al., 2020).

In summary, the quantitative analysis of rod/cone gap junctions from the FIB-SEM data, confirms the presence of gap junctions at the sites identified from the contact analysis of SBF-EM data set e2006 and the confocal analysis of Cx36 labeling. Furthermore, rod/cone gap junctions appear as concentric strings around the post-synaptic opening, as reported in the original freeze fracture data (Raviola and Gilula, 1973; Reale et al., 1978).

### Cone/cone Gap Junctions were not Detected

To evaluate cone/cone coupling, we segmented and partially reconstructed six cone pedicles from dataset FIB-SEM 1. The small size/high resolution of the dataset meant that no cone pedicles were complete. Nevertheless, we found 22 contacts between 6 pairs of cones, 3-4 contacts per pair (Fig. 7, suppl.1A). This suggests that a single cone pedicle could make around 20 contacts with up to six surrounding pedicles given a telodendrial coverage of 1.5.

Although we were able to locate cone/cone contacts, they did not have the typical appearance of a gap junction. While the contacts were direct, without intervening glial processes, there was no clear membrane density. When a nearby rod/cone gap junction was contained in the same frame, it was much more densely stained (Fig. 7, suppl.1B). We examined all 22 cone/cone contacts, but we were unable to identify any typical electron-dense gap junctions in this material. This is surprising given previous reports of cone/cone coupling in mammals.

### Exclusion of rod/rod gap junctions

Previous work has reported the presence of rod/rod gap junctions (Tsukamoto et al., 2001). However, in our FIB-SEM sample of 42 reconstructed rod spherules, we were unable to locate any rod/rod gap junctions. The location and packing density of rod spherules means they are often adjacent, but we found there is usually clear separation between their membranes.

Occasional small contacts between adjacent rod spherules show no membrane density, in contrast to rod/cone gap junctions (Fig. 7, suppl. 1C), and are often distant from the synaptic opening at the base where most Cx36 is clustered in the confocal data. Thus, our FIB-SEM data does not support the presence of rod/rod coupling in the mouse retina. This is consistent with physiological results that show the lack of direct rod/rod coupling (Jin et al., 2020).

## Discussion

We report the reconstruction of photoreceptor terminals and the size and distribution of rod/cone gap junctions in the OPL of the mouse retina. Cone telodendria contact all nearby rod spherules at Cx36 clusters, close to the mouth of the synaptic opening. Confocal microscopy showed there were over 50 Cx36 clusters per cone pedicle with a mean of 2-3 per rod spherule.

Reconstructing an area of 29 cones and 811 rods by SBF-SEM, revealed that cone pedicles, including true blue cones, contacted a mean of 43 rod spherules, reaching > 99% of rods.

Conversely, rod spherules diverge to a mean of 1.9 nearby cones. FIB-SEM, at higher resolution but a smaller volume, confirmed that gap junctions occur at telodendrial contacts with rod spherules. String-like rod/cone gap junctions have a mean length of 480nm with 1.7 gap junctions per rod/cone pair yielding 82 connexons and a theoretical maximum conductance of ∼1200pS, based on morphology. Previous paired recordings showed a mean conductance between a rod/cone pair of 300pS (Jin et al., 2020), suggesting ∼25% of the gap junction channels are open at rest. Rod cone gap junctions are closed by dopamine agonists reducing the conductance to noise levels (∼50pS, ∼5% open probability; Jin et al., 2020). But the mean rod/cone conductance is raised to ∼1000pS in the presence of a dopamine antagonist (Jin et al., 2020), suggesting that the open probability for channels in a rod/cone gap junction may approach 100% and have a dynamic range of ∼20. A summary diagram showing various rod/cone gap junctions is shown in figure 8.

**Figure 8.**
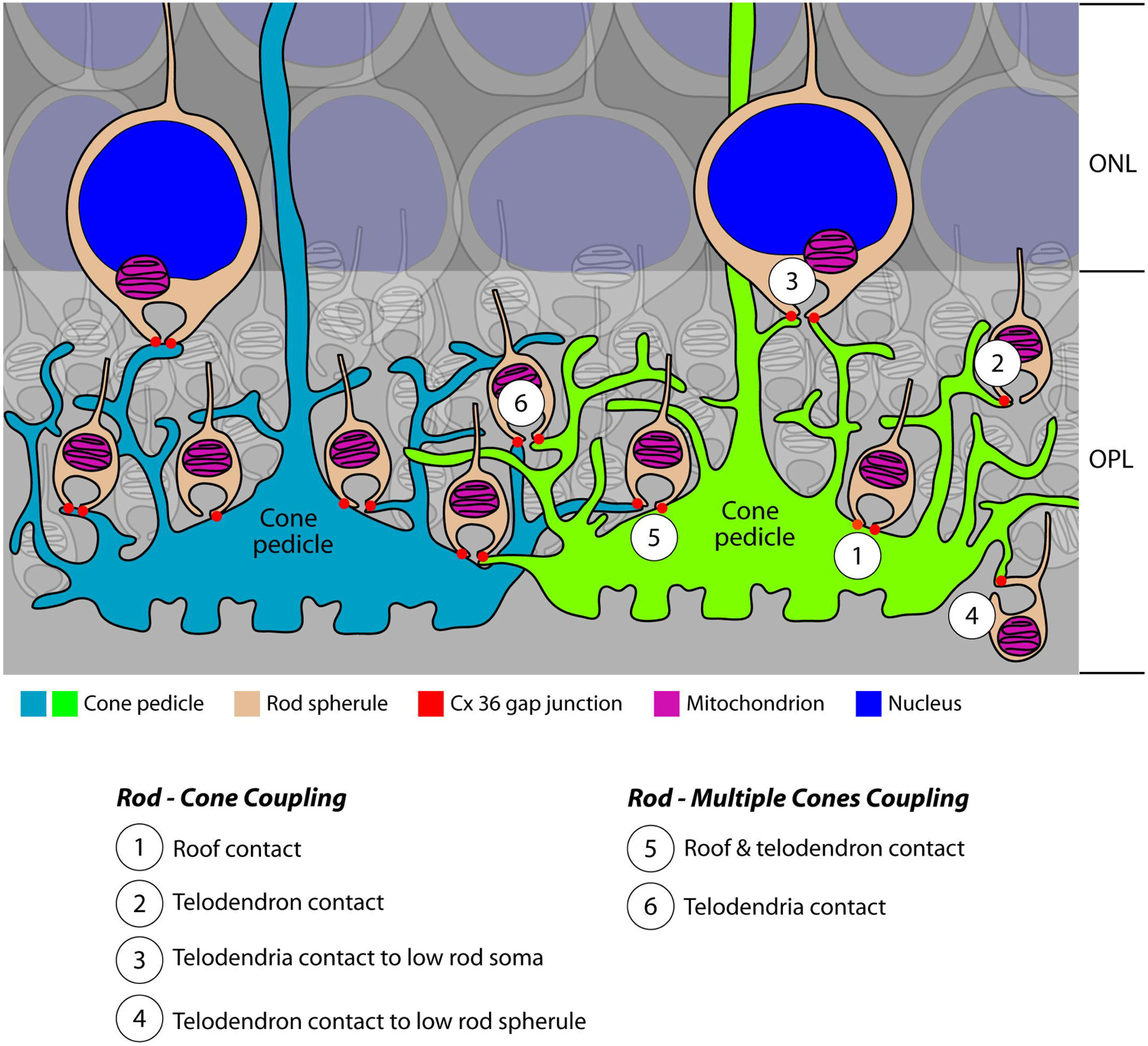
Cartoon Summary of Rod/Cone Gap Junctions in the Mouse Retina A cartoon showing the variety of Cx36 rod/cone gap junctions, including cone telodendrial contacts, cone pedicle roof contacts and inverted rod spherules. The lowest row of rod cell bodies in the ONL do not have rod spherules but still have Cx36 gap junctions with cone telodendria, which reach the upper margin of the OPL. All these structural variants were also found with blue cone pedicles indicating there was no color specificity in rod/cone coupling. Most rod spherules have gap junction contacts with more than one cone. All rod spherules, with very few exceptions (such as areas of low cone density), make gap junctions with nearby cone pedicles. In contrast to the numerous rod/cone gap junctions, we could not detect rod/rod or cone/cone gap junctions in these experiments.

### Identification of gap junctions

In EM material, gap junctions have been difficult to identify due to their size and orientation; unless a gap junction is fortuitously captured in cross section, the membrane density can be difficult to identify. Ideally, there should be enough resolution to reveal the pentalaminar signature of a gap junction, approximately 10nm across (Marc et al., 2018, 1988). Indeed, some authors advocate resampling at high magnification (0.27nm) with goniometric tilt to optimize visibility (Sigulinsky et al., 2020). However, the time investment makes this impractical for every potential gap junction. In our FIB-SEM material, and appropriately fixed SBF-SEM material, the pentalaminar structure of gap junctions cannot be resolved but an area of increased membrane density and chromophilic staining is present where the membranes of a cone telodendron and a rod spherule merge. The identity of the chromophilic material is unknown but presumably it represents the aggregation of connexons and auxiliary proteins at gap junction sites (Lasseigne et al., 2021; Nagy et al., 2018). For rod spherules, this coincides with the location of Cx36 labeling, close to the mouth of the synaptic opening, and so there is a high probability that these sites represent gap junctions.

### Rod/cone Gap Junctions Predominate

Our results confirm the original work from cat retina which showed 48 rod contacts per cone with a divergence of one rod to two cones (Smith et al., 1986), very close to the numbers reported here. In the primate retina, freeze fracture EM showed string-like gap junctions of 400-600 nm in concentric arcs around the rod spherule synaptic opening (Raviola and Gilula, 1973).

The telodendrial contacts described here, have the same curved appearance and coincide with the location of Cx36 on the lower or vitreal surface of each rod spherule. With FIB-SEM, we found string-like rod/cone gap junctions with a mean length of 480nm, a remarkable tribute to the freeze fracture EM work from nearly 50 years ago. Since that time, Cx36 has been identified as the major neuronal connexin and we show that Cx36 labeling signposts rod/cone gap junctions at the confocal level. In summary, the results from mouse, cat, primate and human retinas all indicate that rod/cone gap junctions are a common feature of the mammalian retina (Kántor et al., 2016; O’Brien et al., 2012; Raviola and Gilula, 1973; Smith et al., 1986).

The location of most Cx36 clusters at cone contacts with the base of each rod spherule is consistent with previous EM work showing that gap junctions occur at these sites. In fact, they are quite numerous in this study, with a range of 1-6 Cx36 contacts per rod spherule, comparable with 4-6 cone contacts per rod in the cat retina, forming a ring of contacts around the post-synaptic opening of the rod spherule (Smith et al., 1986). In rod spherules from the mouse retina, two gap junction contacts were described, usually on opposite sides of the synaptic opening, mostly arising from a single cone (Tsukamoto et al., 2001). The detailed reconstructions presented here show a mean of 2.4 (confocal) – 3.2 (FIB-SEM) gap junction contacts per rod spherule and the norm is connections with 1-3 different cones. Our results demonstrate that rod/cone gap junctions account for nearly all gap junctions between photoreceptors.

Previously, some Cx36-GFP labeling was also found in the outer nuclear layer (ONL) (Feigenspan et al., 2004), suggesting that some coupling may occur between somas and/or passing axons. However, we did not observe this pattern with Cx36 antibodies and, in our hands, Cx36 labeling of photoreceptors was restricted to the OPL (Fig. 1). It is possible that the Cx36-GFP construct caused a trafficking defect.

Rod/cone coupling forms the entry to the secondary rod pathway (Kolb, 1977; Smith et al, 1986). It was previously thought that these gap junctions must be closed at night to preserve the amplitude of single photon responses of individual rods, which underlie scotopic sensitivity (Smith et al., 1986). However, recent work indicates that rod/cone coupling is high at night and low in daytime, due to circadian activity and the influence of dopamine (Jin et al., 2016, 2020).

Thus, we have suggested that noise reduction, a consequence of coupling in the photoreceptor network, is a major function of rod/cone gap junctions, in addition to driving the secondary rod pathway (Field et al., 2019; Jin et al., 2020).

### Rods and cones both express Cx36

The small number of Cx36 clusters per rod, compared to more than 50 per cone pedicle, a ratio of approximately 20, may explain early failures to detect rod Cx36 (Bolte et al., 2016; Feigenspan et al., 2004). However, there is strong physiological evidence that Cx36 is required for rod/cone coupling (Asteriti et al., 2017; Ingram et al., 2019; Jin et al., 2020) Cx36 is the most common neuronal connexin but it does not pair with other connexins, making only homotypic gap junctions (Koval et al., 2014; Nagy et al., 2018; Teubner et al., 2000). We have recently shown that both rods and cones express Cx36 and that no other connexins are present in either rod or cones (Jin et al., 2020). In addition, Cx36 is required on both sides to form a rod/cone gap junction (Jin et al., 2020; Miller et al., 2017). Thus, the simplest case holds, that both rods and cones express Cx36 and rod/cone gap junctions are heterologous but homotypic (meaning between different cell types but both expressing the same connexons).

### Blue cones are also coupled to rods

Blue cones, identified in confocal work by the presence of S-cone opsin, and in SBF-SEM by their connections with blue cone bipolar cells (Behrens et al., 2016; Nadal-Nicolás et al., 2020), also made telodendrial contacts at Cx36 clusters with all nearby rod spherules (Fig. 4). In the cone networks of primate and ground squirrel retina, there is evidence that blue cones are not coupled to neighboring red/green or green cones (Hornstein et al., 2004; Li and DeVries, 2004; O’Brien et al., 2012), perhaps to maintain spectral discrimination (Hsu et al., 2000). However, we find no evidence for color specificity in rod/cone coupling. In fact, a single rod spherule may be coupled to both blue and green cones (Fig. 5, supplement 5). Thus, rod input to blue cone bipolar cells and downstream circuits is predicted via the secondary rod pathway, in addition to the previously reported primary rod pathway inputs from AII amacrine cells to blue cone bipolar cells (Field et al., 2009; Whitaker et al., 2021)

### Cx36 numbers for rods and cones are consistent

We have gathered data for both rods and cones independently. Because they are linked by rod/cone gap junctions, a book-keeping exercise should serve as a check on the data. From individual EGFP-labeled cone pedicles, we estimate that each cone pedicle has 51 ± 8.9 (mean ± SD, n=18) Cx36 clusters, presumed to represent gap junctions, with nearby rod spherules. From the rod perspective, 43 rods contact each cone with 2.4 Cx36 clusters per rod. But some of these potential gap junctions are made with neighboring cones. To correct for this, we divide by the rod divergence, 1.9. Thus 43 x 2.4/1.9 = 54, the number of Cx36 clusters at rod contacts with a single cone pedicle. This is in close agreement with the number of Cx36 gap junctions counted in confocal reconstructions of individual cone pedicles. These numbers are also close to previous calculations of ∼45 Cx36 clusters/cone pedicle, based on the density of Cx36 puncta and the number of cone pedicles from wholemount retina (Jin et al., 2020).

We calculated the number of gap junctions per rod by two methods, a confocal analysis of Cx36 labeled points, and using the FIB-SEM data. Analyzing the FIB-SEM data gave 3.2 gap junctions per rod spherule. In the confocal analysis of Cx36 elements, there were 2.4 gap junctions per rod spherule, 25% less. While these numbers are similar, the EM data gives a higher value. This might be expected because small elements could be missed and two close together could merge and be counted as one in the lower resolution confocal data (Sigulinsky et al., 2020). Clearly, the smallest Cx36 elements may be less than the confocal detection limit, (different from the resolution limit), but these data suggest that we accounted for at least 75% of the photoreceptor gap junctions by confocal microscopy.

### Cone/Cone Contacts were not Detected

We were unable to confirm the presence of cone/cone gap junctions in the mouse retina, despite locating more than 20 examples of cone/cone contacts. In each case, there was no indication of the typical membrane density or chromophilic staining at cone/cone contacts. Nearby rod/cone contacts provided prominent control gap junctions for comparison (Fig. 7 suppl. 1B). This result is surprising, given previous reports of cone/cone coupling in several mammalian species, including mouse (DeVries et al., 2002; Kolb, 1977; Smith et al., 1986; Tsukamoto et al., 2001). However, close contacts without gap junctions are not unprecedented. In rabbit retina, many bipolar cells are coupled by gap junctions, but some are not, despite contacts and the presence of gap junctions elsewhere in the same cell (Sigulinsky et al., 2020). Gap junctions do not determine the specificity of neuronal connections.

However, electrophysiological studies suggest there is weak cone to cone coupling in the mouse retina, which may indicate that cone/cone gap junctions are smaller than rod/cone gap junctions. Cone/cone coupling persists in the rod-specific Cx36 KO, suggesting it is direct (Jin et al., 2020). In the rod Cx36 KO, which should reveal cone/cone coupling, there was some residual Cx36 labeling, associated with cone telodendria, which was significantly different from the background noise and may represent a small amount of cone/cone coupling, estimated as 2 gap junctions/cone (Jin et al., 2020).

Cone to cone coupling has been reported in other mammals but there may be species variation. In the ground squirrel, a cone dominated retina, the low number of rods with the resulting adjacency of the cones may promote cone/cone coupling (DeVries et al., 2002; Li, 2020; Li et al., 2010). In the central primate retina, cones are densely packed and often adjacent, which is not the case in mouse retina. In peripheral primate retina, cones are more widely spaced, and they are connected by a sparse array of telodendria, which seem to target neighboring cones making the pattern of Cx36 labelled gap junctions very obvious, in addition to numerous rod/cone gap junctions (O’Brien et al., 2012). Finally, in the rod-less, cone-only mouse, there is a large increase in Cx36 labeling in the OPL suggesting that cone/cone coupling occurs, at least under these circumstances (Dang et al., 2004). The evidence for photoreceptor coupling, including weak cone/cone coupling, is summarized in Appendix 1.

### No rod/rod gap junctions

Previous work in the mouse retina has suggested that rods form a coupled network based on the presence of small rod/rod gap junctions (Tsukamoto et al., 2001). Most of the rod spherules were reported to make gap junction contacts which were characterized as small and with no membrane density (Tsukamoto et al., 2001). Furthermore, some of the rod/rod contact sites were between the spherules and passing rod axons, a location where there is no Cx36 labeling.

We have searched diligently for rod/rod gap junctions, but we have been unable to confirm their presence. Not only is Cx36 labeling restricted to the base of the rod spherule, around the synaptic opening, but, in the FIB-SEM material, this was almost the only location where we found gap junctions. We mapped the gap junctions from 42 rod spherules and their distribution was in close agreement with the distribution of Cx36 labeling and cone telodendrial contacts. We could not trace every process making a gap junction contact but in 112/112 cases, we could identify the process as a cone telodendron. Even the small number of outlying gap junctions distant from the synaptic opening were traced to cones.

We did find small contacts between adjacent rod spherules (Fig. 7, suppl. 1C) but there was no membrane density indicating the presence of a gap junction and, in this equatorial position, around the midline where the rod spherules are closely packed, there was no Cx36. We are aware of the difficulty in proving the absence of a particular structure but, based on the evidence presented here, we conclude that there are no rod/rod gap junctions in the mouse retina and this may be a common feature of the mammalian retina.

In support of this conclusion, apparent rod/rod coupling was eliminated in the cone-specific Cx36 KO, indicating that physiological evidence of rod/rod coupling in wild type retina is actually due to indirect rod/cone/rod coupling via the network (Jin et al., 2020). In the cone-specific Cx36 KO, which should reveal rod/rod coupling, Cx36 labeling of photoreceptors was essentially eliminated, providing no evidence for residual rod/rod coupling (Jin et al., 2020). The rod-specific Cx36 KO produced a similar result but a few remaining Cx36 clusters were attributed to cone/cone coupling. The loss of most Cx36 coupling in either the rod- or cone-specific KO indicates that rod/cone coupling accounts for the majority of photoreceptor gap junctions and provides no support for direct rod/rod coupling. We must emphasize that the weight of the available evidence suggests that there is no rod/rod coupling in the mouse retina.

The comparative evidence for photoreceptor coupling, including the lack of rod/rod coupling, is summarized in Appendix 1.

### Calculations of Size and Open Probability for Rod/Cone Gap Junctions

Gap junctions are common building blocks of neural circuits throughout the CNS, but there is much that we do not know. For example, in most cases, we do not know their size, the number of connexon channels, their conductance, and perhaps most importantly, their dynamic range or plasticity. The size of the conductance through a gap junction depends not only on the number of connexons and the unitary conductance, but also on the channel activity, usually characterized as the open probability. These are the fundamental properties required to understand the function of gap junctions in neuronal microcircuits. Here, we have an opportunity to estimate these variables.

There is great variability in gap junction morphology, from strings to crystalline plaques (Kamasawa et al., 2006). Rod/cone gap junctions appear as low-density strings of single particles in freeze fracture EM (Raviola and Gilula, 1973; Reale et al., 1978) and this is consistent with the uniform width of rod/cone gap junctions from our 3D-EM reconstructions, as well as the low intensity of Cx36 labeling compared to the plaque type gap junctions of the inner retina (Kamasawa et al., 2006).

We calculated the number of connexon channels from the mean length of an FIB-SEM gap junction ∼ 480nm, yielding 48 channels per rod/cone gap junction, based on the 10nm spacing of connexons in a string (Kamasawa et al., 2006; Szoboszlay et al., 2016), in close agreement with 40-50 particles/string in freeze fracture (Raviola and Gilula, 1973). We estimate there are 3.2 gap junctions per rod, but these are shared with several nearby cones. Dividing by the rod to cone divergence (the number of cones contacted by each rod), 1.9, gives 1.7 gap junctions or a mean of 82 connexons between an average rod/cone pair. For a unitary conductance of 15 pS per Cx36 channel (Srinivas et al., 1999), we can calculate the mean maximal conductance between a rod and a cone as ∼ 1,200 pS (82 connexons x 15 pS/connexon). It is important to note that this theoretical maximum, if all gap junction channels between a rod/cone pair are in an open state, was derived from our morphological data.

Using this detailed structural information, combined with our previous conductance measurements from rod/cone paired recordings, we can calculate the properties of rod/cone gap junctions. Our recently published work shows a resting value ∼ 300 pS for the rod/cone transjunctional conductance (Jin et al., 2020). Compared to the theoretical maximum above, this indicates a resting open probability of 25%. Notably, this is much higher than previous estimates of around 1% (Connors, 2017; Marandykina et al., 2013). But these earlier values may be low because the size of the gap junctions was estimated by immunofluorescence, which provides a high number for gap junction area and connexon number, with a correspondingly low number for the open probability (Kamasawa et al., 2006; Szoboszlay et al., 2016). More recently, when the number of connexons was accurately measured by freeze fracture EM, the open probability of cerebellar gap junctions was calculated as 18% (Szoboszlay et al., 2016), close to the value derived here.

### Dynamic Range of Rod/Cone Gap Junctions

Rod/cone coupling is modulated by dopamine (Ribelayga et al., 2008; Jin et al., 2020). In our previous studies, we showed that the dynamic range of rod/cone coupling, driven by D2 dopaminergic agonists and antagonists, varied from a noise level ∼50pS in the closed state to a mean maximum of 1070pS (Jin et al., 2020). Compared to the theoretical maximum above, these values translate to a minimum open probability of 4% and a maximum of 89% of available Cx36 channels. This is a very surprising result because previous measures of open probability have produced such low estimates (Connors, 2017), but we believe it is consistent with our data. For the first time, it suggests that the range of gap junctions can be modulated from close to 0 to nearly 100% of available Cx36 channels. In other words, all of the channels in these small rod/cone gap junctions are switchable and may contribute to their plasticity.

The diffusion coefficient through Cx36 gap junctions, a proxy for conductance, was correlated with the phosphorylation of Cx36 as measured using phospho-Cx36 antibodies (Kothmann et al., 2009; O’Brien, 2014). Pharmacological manipulation using dopamine agonists or antagonists produced phosphorylation-driven changes in tracer coupling that encompassed a 20-fold range of diffusion coefficients, producing a large dynamic range for gap junction plasticity. In the present experiments, the ratio of minimum to maximum open channels, from 4% − 89% or 50-1000pS, was also approximately 20. This is very similar to the dynamic range derived from tracer coupling studies with phospho-specific Cx36 antibodies. While these calculations are necessarily approximate, they support the concept that gap junctions provide a versatile component, offering a large range of plasticity in neural circuits. In the retina, rod/cone coupling may reduce transduction noise in the photoreceptor network and the modulation of rod/cone gap junctions also provides a switchable entry to the secondary rod pathway, which varies with light intensity and/or the circadian cycle (Bloomfield and Völgyi, 2009; Field et al., 2019; Jin et al., 2020; Jin and Ribelayga, 2016).

## Methods

### Animals

All animal procedures were reviewed and approved by the Animal Welfare Committee at the University of Texas Health Science Center at Houston or by our collaborators’ local Institutional Animal Care and Use Committees. C57BL/6J (stock no. 000664) mice were purchased from the Jackson laboratories. We used mice 2 to 6 months of age of either sex. Animals were housed under standard laboratory conditions, including a 12-hour light/12-hour dark cycle.

### Antibodies and immunocytochemistry

Mice were anesthetized by intraperitoneal injection of a ketamine/xylazine mix solution (100/10 mg/kg) before being euthanized by cervical dislocation. Eyes were rapidly collected, hemisected, the vitreous was removed and the resulting eyecup was placed in 4% paraformaldehyde in phosphate-buffered saline (PBS) at room temperature for 1 to 2 hours.

Vibratome sections (Leica VT 1000S) or whole-mounted retinas were reacted with a cocktail of antibodies (Table 1), according to procedures described previously (Li et al., 2013; O’Brien et al., 2012). Briefly, sections were washed and blocked in 3% donkey serum/0.3% Triton X-100 (in PBS) for 2 hours (overnight for whole mounts) and incubated overnight at room temperature with a cocktail of primary antibody(ies) in 1% donkey serum/0.3% Triton X-100 (in PBS) (7 days for whole mounts). Tissues were processed free-floating on an oscillating platform at 1Hz.

Following incubation with the primary antibody, sections were rinsed in PBS (6×, 20 min) and reacted with a secondary antibody(ies) for 2 hours (overnight for whole mounts) at room temperature in the dark. Donkey Alexa Fluor–, Cy3-, or DyLight-conjugated secondary antibodies were purchased from Jackson ImmunoResearch Laboratories Inc. (West Grove, PA) and used at 1:600 dilution. Last, sections or whole mounts were covered with mounting medium, 6-diamidino-2-phenylindole) (100 μ mounting medium to stain the nuclei.

### Blue cone opsin-Venus mouse line

A bacteria artificial chromosome (BAC) clone (bMQ-440P15) from a SV129 genomic library (bMQ), encompassing the entire mouse Opn1sw locus, was used to create the Opn1sw_Venus BAC transgenic construct through ET recombination as described elsewhere (PMID: 9771703). Briefly, a Venus-bGH PolyA-NeoR (flanked by FRT sites) cassette was integrated into the Opn1sw locus on the BAC clone replacing coding region of exon 1 to generate the Venus-bGH-Neo knockin version of the BAC clone. Recombination was verified by PCR reactions specifically designed to detect recombination junctions by ET recombination (or homologous recombination). This BAC clone carrying the reporter cassette was further trimmed, through ET recombination, to retain a 15 kb fragment including upstream regulatory sequence (10kb upstream of the coding region), the KI cassette and the ∼2 kb sequence downstream of exon 2 of the Opn1sw gene. The intended transgenic construct was verified by overlapping PCR and Sanger sequencing. Transgenic F0 founders were created by microinjecting the transgenic construct (1ng/µl) into fertilized eggs from C57bl6/j donors followed by genomic PCR screening amplifying the reporter insert.

### Confocal Microscopy Image Acquisition

A Zeiss LSM-800 confocal microscope with Airyscan was used for 4 channel imaging of retinal whole mounts and sections with a 63x (N.A. 1.4) oil-immersion objectives. Images with 30 - 40 nm pixel size were acquired in series of 0.15 μ processed by Zen software (Zeiss) to yield super-resolution images. We measured the full width at half maximum of the point spread function for this instrument as 170nm using 0.1 μ beads (Molecular Probes). Figures were presented as short stacks of 2 - 6 images. Images were processed in ImageJ, Imaris (Oxford Instruments) or Photoshop (Adobe Systems Inc) for contrast enhancement and further analysis.

### Quantitative Analysis of Cx36 clusters

Single cones: retinal sections from OPN4-EGFP mice were used because only a few cones were EGFP positive and it was possible to image well-isolated single cone pedicles. Airyscan images were loaded in Imaris software and the colocalization between the cone (EGFP) and Cx36 (Cy3) was analyzed. Neighboring colocalized voxels were grouped as a 3D object by surface rendering and the number of extracted 3D objects was measured.

Rod spherules: an antibody against the synaptic vesicular glutamate transporter (vGlut1) was used to label rod spherules. It does not stain the cell membrane and thus underfills rod spherules (Quraishi et al., 2007). Some sections were also stained for Plasma Membrane Calcium ATPase (PMCA) which stains the rod spherule membrane (Johnson et al., 2007), encircling the vGlut1 labeling (Fig. 5, suppl. 3A). Due to lack of clear colocalization between Cx36 and the rod marker (vGlut1), the number of Cx36 clusters per rod was counted manually. Some rod spherules were excluded from analysis if they were too close to link a Cx36 cluster to a specific rod. A total of 116 well-isolated rod spherules in 3 retinal sections were chosen to analyze for the number of Cx36 clusters. To analyze the position, a mini-stack of 6 x 0.15 μm optical sections from 18 rod spherules were extracted, aligned and averaged (ImageJ). A spline curve 8 pixels (272nm) wide was applied to the perimeter and the intensity profile for Cx36 was plotted linearly around the synaptic opening.

### Connectivity analysis, SBF-SEM (e2006)

A region in the OPL from a publicly available serial block-face scanning EM (SBF-SEM) dataset e2006, voxel size 16.5 x 16.5 x 25 nm, (Behrens et al., 2016; Helmstaedter et al., 2013), including 164 cone pedicles was analyzed using KNOSSOS software (Helmstaedter et al., 2011) m in the OPL was chosen to analyze, in which we identified 29 cone pedicles and 811 rod spherules. Because there are some areas of relatively low cone density, an additional area including 6 widely spaced cone pedicles was also analyzed. The whole e2006 dataset included a total of 6 blue cones (Behrens et al., 2016) but one was located at the edge of the dataset and was truncated. Thus, a total of 5 blue cones were analyzed.

Skeletons: we traced the telodendria from all 29 cone pedicles by skeletonization with KNOSSOS. All rod spherules contacted by a specific cone were identified. We used the membrane fusion between cone pedicle/telodendria and rod spherule to indicate rod/cone contact. The contact information was stored as an annotation directory in KNOSSOS and further analyzed using Excel (Microsoft) and Origin (OriginLab Corp). μm (OPL plus a part of ONL) × 27 μ m was extracted from the e2006 dataset to segment using Microscopy Image Browser (Belevich et al., 2016). The segmented voxel data were loaded into Imaris to create a 3D reconstruction by surface rendering. Images were constructed in Imaris by rotating and adjusting transparency as required.

### SBF-SEM, Singer dataset (eel001)

An excised retina was fixed for one hour at room temperature with 2% glutaraldehyde in 0.15 M cacodylate buffer, washed in three changes of the same buffer, and postfixed with 1% osmium tetroxide in 0.15 M cacodylate containing 1.5% potassium ferrocyanide. A wash in three changes of distilled water followed the reduced osmium fixation and preceded an en bloc fix in 2% aqueous uranyl acetate. Dehydration in a graded series of ethanol (35% to 100%) and infiltration in a propylene oxide:epoxy resin series was followed by embedding and polymerization in epoxy resin. Imaging by SBF-SEM was performed under contract with Renovo Neural (Cleveland, OH) using a FEI Teneo Volume Scope. Beam current was 50 pA and the face was imaged with 7 x 7 x 40 nm resolution. Each face image spanned the width of the retina from the first layer of photoreceptor cell bodies in the ONL to the ganglion cell bodies in the GCL and comprised two stitched fields of 8192 x 8192 pixels for a dimension of 8192 x 16384 pixels or 57.34 x ∼114.68 µm. 1649 slices in total were acquired, making the z-depth traversed = 65.96 µm. Image stacks were viewed using Knossos and processed in Photoshop.

### FIB-SEM sample preparation

Mice were euthanized and enucleated; retinas were quickly dissected and placed in 3% paraformaldehyde and 1% glutaraldehyde for 30 minutes, then processed for electron microscopy using the Dresden protocol (Paridaen et al., 2013). The resulting resin blocks were trimmed to contain the OPL within 25μm of the top-edge of the block, cut and mounted to a 45° μ pre-tilted stub, and coated with 8nm of carbon. Three-dimensional data were acquired using the Helios G3 NanoLab DualBeam FIB-SEM. In brief, a focused beam of gallium atoms ablated 4nm off the surface of the sample (FIB conditions: 30keV accelerating voltage, 790pA beam current), comprising the scanning electron microscope (SEM) imaging area, with a field of view around 25μm. The freshly ablated surface was then imaged by backscattered electrons using the In-Column Detector (SEM conditions: 400 pA beam current, 3KeV accelerating voltage, 4nm per pixel, and 3μs dwell time; image resolution: 6144 x 4086, 4nm isotropic). Around 1,149 (FIB-SEM 1 dataset; 1,342 for FIB-SEM 2 dataset) ablation and imaging cycles were run over a three day period, resulting in a sample depth of roughly 4.6μ using the Linear Stack Alignment with SIFT algorithm (Fiji) and cropped to remove artifacts arising from the alignment.

### Gap junction position and size analysis from FIB SEM

Individual rod spherules were cropped from the datasets and segmented for 3D reconstruction as described above for the e2006 dataset. Gap junctions were identified by a darkly stained area of merged membranes. The distance from a gap junction to the synaptic opening of a rod spherule was calculated using the oblique slicer tool in Imaris to align the center of the mitochondria and the postsynaptic opening in a single plane. Gap junction length was visualized using the oblique slicer to create an en face view. Some gap junctions showed belt-like structures with curvature and torsion. Due to the complexity of the structure, the size of these gap junctions was estimated as a simplified rectangular shape by measuring the length and average width of the belt. The acquired data were analyzed in Excel and Origin.

**Table 1:**
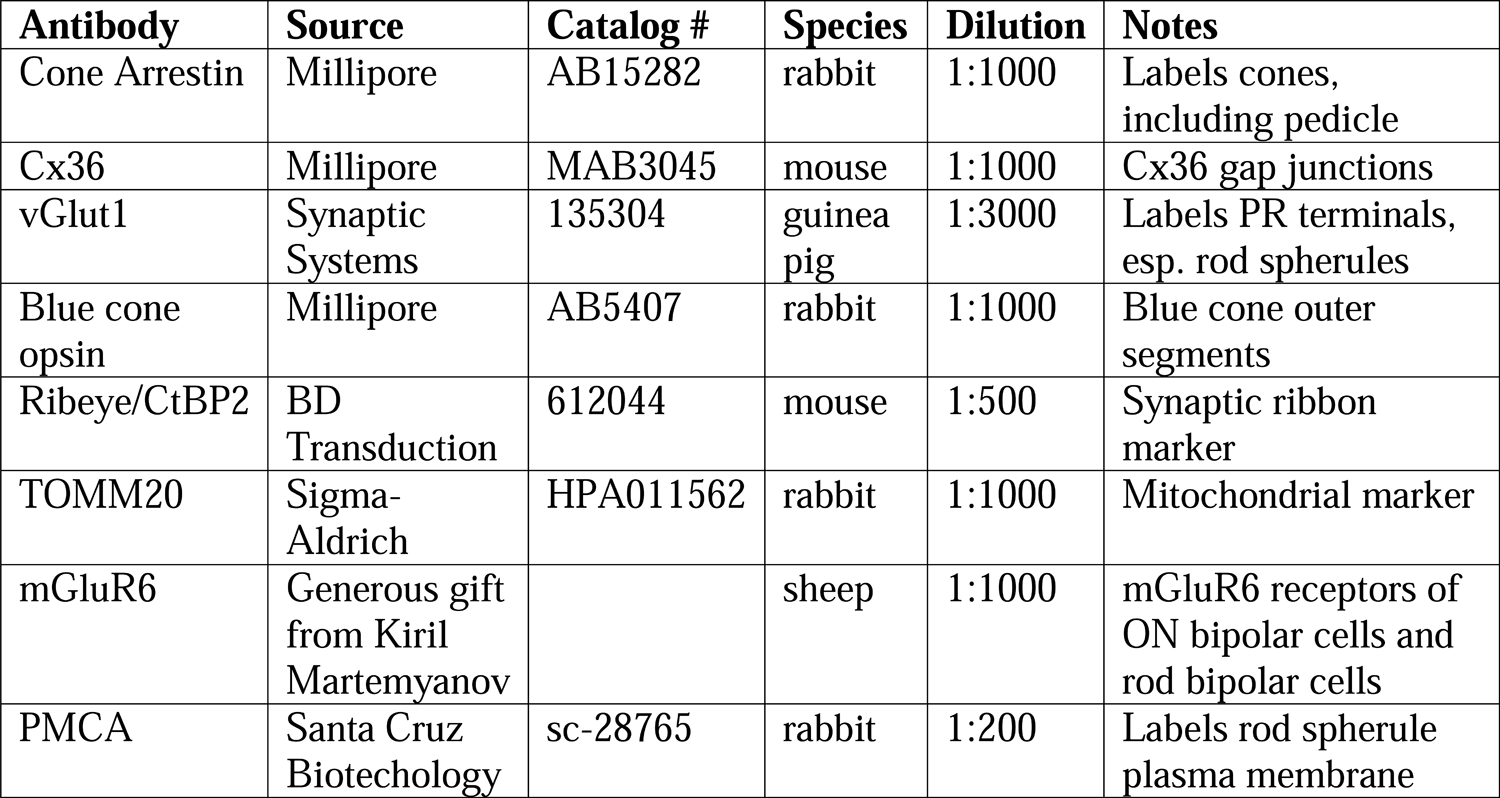
Antibodies

| **Antibody** | **Source** | **Catalog #** | **Species** | **Dilution** | **Notes** |
| --- | --- | --- | --- | --- | --- |
| Cone Arrestin | Millipore | AB15282 | rabbit | 1:1000 | Labels cones, including pedicle |
| Cx36 | Millipore | MAB3045 | mouse | 1:1000 | Cx36 gap junctions |
| vGlut1 | Synaptic Systems | 135304 | guinea pig | 1:3000 | Labels PR terminals, esp. rod spherules |
| Blue cone opsin | Millipore | AB5407 | rabbit | 1:1000 | Blue cone outer segments |
| Ribeye/CtBP2 | BD Transduction | 612044 | mouse | 1:500 | Synaptic ribbon marker |
| TOMM20 | Sigma-Aldrich | HPA011562 | rabbit | 1:1000 | Mitochondrial marker |
| mGluR6 | Generous gift from Kiril Martemyanov |  | sheep | 1:1000 | mGluR6 receptors of ON bipolar cells and rod bipolar cells |
| PMCA | Santa Cruz Biotechnology | sc-28765 | rabbit | 1:200 | Labels rod spherule plasma membrane |

## Appendix 1

### Comparison of Photoreceptor Gap Junctions

For comparison, we have summarized the evidence for rod/cone, cone/cone, and rod/rod gap junctions in the mouse retina.

### Rod/cone

1. Cones contact all rods.
2. Cx36 labeling at the base of each rod spherule; no other connexins expressed by photoreceptors (Jin et al., 2020).
3. Cx36 labeling dramatically reduced in either rod or cone specific Cx36 XOs, consistent with the predominance of rod/cone gap junctions and the requirement for Cx36 on both sides of a rod/cone gap junction (Jin et al., 2020).
4. FIB-SEM shows gap junctions at the base of each rod spherule, coincident with cone contacts and Cx36 immunoreactivity.
5. Paired recordings show a large conductance between rod/cone pairs (resting state, dark-adapted), mean 300pS (Jin et al., 2020).
6. Rod/cone coupling abolished in either the rod- or cone-specific Cx36 XO (Jin et al., 2020).
7. Rod signals transmitted to cones via rod/cone gap junctions (Jin et al., 2020).
8. Rod/cone gap junctions are common in several mammalian species (Smith et al, 1986; O’Brien et al., 2012).

We conclude all rods are coupled to cones.

### Cone/cone

1. Some cone/cone contacts, mostly between telodendria.
2. While Cx36 is colocalized with the cone telodendrial network, most Cx36 clusters are rod/cone GJs. We were unable to identify cone/cone Cx36 clusters in the mouse retina.
3. FIB-SEM shows some contacts between adjacent cones but no membrane density at contact points.
4. Paired recordings show weak cone/cone coupling, a fraction of rod/cone coupling, ∼60pS in mouse (Jin et al., 2020), but ∼250pS in ground squirrel where Cx36 between cones is prominent (DeVries et al., 2002).
5. Cone/cone coupling was preserved in the rod Cx36 XO, suggesting cone/cone coupling is direct.
6. In the rod XO, which should reveal cone/cone coupling, a few Cx36 clusters left, significantly different from cone Cx36 XO or pan Cx36 XO, (Jin et al., 2020).
7. Cone/cone coupling reported in other species, such as ground squirrel and primate,
8. In the rod-less, cone-only mouse, there is a large increase in Cx36 labeling in the OPL (Dang et al., 2004).

The evidence for cone/cone coupling is weak for the mouse retina. Despite the fact that we were unable to identify cone/cone gap junctions, there is physiological evidence for weak cone/cone coupling (Jin et al., 2020). While the presence of cone/cone coupling in the mouse is equivocal, there is strong evidence for cone/cone coupling in other species such as primate and ground squirrel. (DeVries et al., 2002; O’Brien et al., 2012).

### Rod/rod

1. Few rod/rod contacts despite spherule packing in the OPL
2. No Cx36 labeling at rod/rod contacts
3. In the cone-specific Cx36 XO, which should reveal rod/rod coupling, there was no remaining Cx36 labeling in the OPL above a very low background (Jin et al., 2020).
4. FIB-SEM of rod/rod contacts shows no dense staining typical of a gap junction
5. Paired recordings show apparent rod/rod coupling was abolished in the cone Cx36 XO, indicating indirect or network coupling, rod/cone/rod, (Jin et al., 2020).
6. Cx36 in the ONL is very weak, indistinguishable from non-specific background labeling. If any immunolabeling represents gap junctions, as opposed to non-specific noise, these very low numbers are not enough to account for coupling between the multitude of rods

We conclude there is no direct rod/rod coupling in the mouse retina.

In summary, the evidence for the predominant role of rod/cone gap junctions is strong and straightforward. In contrast, we found no evidence for rod/rod gap junctions and the distribution of Cx36 does not match the location of rod/rod contacts. We were also unable to detect cone/cone gap junctions by immunofluorescence or EM, but there is at least some supporting physiological evidence in favor of cone/cone gap junctions. It may seem unsatisfactory that we were unable to detect rod/rod or cone/cone gap junctions, but it does not change the main result: that rod/cone coupling accounts for the vast majority of photoreceptor gap junctions.

## Appendix 2

## Supplementary Figure Legends

**Figure 1, supplement 1** The distribution of blue cone opsin in the mouse retina Wholemount view of whole blue cone opsin Venus mouse retina shows most cones in ventral retina express blue cone opsin. In the dorsal retina, there is a sparse mosaic of true blue cones.

**Figure 1, supplement 2** Blue cone pedicles have telodendria bearing Cx36 clusters. **A.** Dorsal retina of a blue cone opsin Venus mouse contains a sparse set of true-blue cones (cyan) among the majority of green cones labeled for cone arrestin (green). **B.** Zoom in on one blue cone pedicle shows Cx36-labeled (red) telodendria. Surrounding green cone pedicles labeled for cone arrestin (green). Large red structures are non-specifically labeled blood vessels. **C.** Outer segments showing a sparse set of cones labeled with an antibody against blue cone opsin (cyan). Background of green cones labeled for cone arrestin (green). **D.** Following the cone axons through a series of confocal images allows the blue cone pedicles to be identified (asterisks), see supplementary movie 3 from Fig. 1, supplement 2. **E.** Enlarged view of a single blue cone pedicle (asterisk) with Cx36 (red) labeled telodendria, comparable to the surrounding green cones labeled for cone arrestin (green)

**Figure 4, supplement 1** Nearly All Rods receive Cone Contacts **A.** Map showing telodendria of all 29 cone pedicles (Fig. 3A, box 1), each individually colored, dots mark all rod spherules in contact, same color as cone if contacts are exclusive to one cone, black, contact with two cones, grey, contact with 3 cones, light grey no contacts, mostly outside the field of all 29 cones. Within the field of 29 cones, there were 811 rod spherules of which 808 (99.6%) received cone contacts. 3/811 (0.4%) rod spherules received no cone contacts. **B.** Removing cone pedicles and rod spherules in contact with the central 13 cone pedicles, leaves a hole with no remaining rod spherules (0/361) inside an annulus of rod spherules outside the field of the central 13 cones. The telodendrial field of cone 5 (green, axon removed) is shown for comparison. **C.** Polygons outline the telodendrial fields of 6 cones in a sparse region (Fig.3A, box 2). In the center, there is a rare hole in the cone coverage, almost like a cone is missing. **D.** Removing cones and plotting rod spherules, (color coded by cone for exclusive contacts, black, two cones, dark grey, three cones, no contacts, light grey) reveals 8/175 (4.6%) of rod spherules in the central area with no cone contacts. This may represent a local upper limit of rods without cone contacts, since we deliberately chose a sparse area.

**Figure 4, supplement 2** Rod Connectivity Map for 29 Cone Pedicles **A.** Full connectivity map for all 29 cone pedicles that were reconstructed, central 13 clear boxes, outer 16, shaded boxes. Each square represents one cone, containing the identity with the total number of rod contacts above, 43 ± 5.4 (mean ± SD, n=29) and the number of exclusive contacts below, 7.2 ± 3.4 (mean ± SD, n=13, central cones only; outer cone numbers for exclusive contacts are high due to edge effects). Circles show number of rod spherules shared between cone pedicles connected by lines. Most cone pedicles share rod spherules with all neighboring cones. Cones 5 (green), 3 (cyan) & 2 (blue) were segmented and reconstructed in 3D. Cone 2 is a blue cone (Behrens et al., 2016). **B.** The number of rod spherules shared between any two adjacent cone pairs, 6.2 ± 4.7 (mean ± SD, n=79 pairs among all 29 cones, black), central 13 cones (red) (5.9 ± 4.4, mean ± SD, n=61 pairs); blue cones, 6.6 ± 6.3 (mean ± SD, n=16 pairs from 5 blue cones). Box shows quartiles, mean (circle), median (bold line), SD (whisker), min/max (x). *79 pairs from the 29 cones include GC-GC and BC-GC pairs while blue cone data shows BC-GC pairs only. Note that there are no BC-BC pairs. GC, green cone, BC, blue cone. Table 2, Appendix 2, additionally shows statistics for only GC-GC pairs, 6.3 ± 4.1 (mean ± SD, n=69 pairs from 31 green cones), (27 GC of the 29 cones and 4 GC in sparse density area).

**Figure 5, supplement 1** Confocal Microscopy, Roof Contacts **A.** Cx36 clusters (red) occur where 3 rod spherules (vGlut1, blue) make contact (arrows) with the upper surface or roof of a single EGFP labelled cone (green). Each of these probable gap junctions is close to the opening of a rod spherule post-synaptic compartment.

**Figure 5, supplement 2** Low Rods A. Confocal microscopy, 3 rod somas, labelled with DAPI, (grey, asterisks) in the lowest row of the ONL adjacent to the upper margin of the OPL, have no axons or spherules. Instead, the synaptic machinery, stained with vGlut1 (blue), is contained in a crescent at the base (Li et al., 2016). Cone telodendria, stained for cone arrestin (green) contact each of these rods with Cx36 clusters (arrows, red) at the contact points, close to the mouth of the synaptic opening. **B.** Segmentation from e2006, single sections; left, a rod soma from the lowest row of the ONL contains post-synaptic inclusions at the base and receives contact from a cone telodendron (green) close to the post-synaptic opening (25nm x 7 sections = 175 nm away); right, a similar example, a low rod soma, (brown) with a post-synaptic compartment at the base receives contacts from two different cones (green and cyan) (16.5 nm x 5 sections = 82.5 nm to the post-synaptic opening). **A. C.** 3D reconstruction of the 5 rods in the lowest row of the ONL above cones 5 & 3, (a subset of the rods in Fig. 5G) shows they sit on a nest of up-reaching cone telodendria. For comparison, a rod spherule (blue) is included, far left, arrowhead. Arrows, axon ascending from rod spherule and cone 5 axon (green). **B. D.** The same 5 low rods rotated to view the cone telodendrial contact pads (yellow or red) around the mouth of the post-synaptic compartment on the lower surface of each rod. Arrows show axons. Inset for size comparison, a reconstructed rod spherule.

**Figure 5, supplement 3** Low Rod Spherules A. Confocal microscopy, OPL, a cone pedicle and telodendria stained for cone arrestin (green) and a cluster of rod spherules, stained for vGlut1(blue) and outlined for Plasma Membrane Calcium ATPase (PMCA) (yellow). Cx36 clusters (red) occur at cone contacts with the base of each rod spherule, close to the synaptic opening (arrowheads). One rod spherule is very low and protrudes below the level of the cone pedicle. This rod spherule is rotated horizontally so the post-synaptic opening is adjacent to the cone pedicle and there is a Cx36 cluster at this contact site (arrow). Thus, the rule is that rod/cone contacts occur close to the rod spherule post-synaptic opening, whatever the orientation. **B.** Segmentation, e2006 dataset, a low rod spherule (pink) is inverted to receive a cone contact (green) close to the synaptic opening (arrowhead), 25 nm x 24 frames = 600 nm away. **C.** An inverted rod spherule (orange), adjacent to a cone pedicle (green) but with clear membrane separation and no contact, receives cone telodendrial contacts (cyan) near the synaptic opening, (25 nm x 12 frames = 300 nm away) which in this case is uppermost. **D.** 3D reconstruction e2006, a low rod spherule (purple) is inverted and receives a cone contact (green, transparent), close to the post-synaptic mouth. Arrows indicate cone axon and rod axon, ascending to rod soma. **E.** Detail showing the contact pad (red) curved around the post-synaptic opening.

**Figure 5, supplement 4** Blue Cones, Segmentation, e2006 **A.** Detail of cone pedicle skeletons from Fig. 4 with the field of cone 2 outlined. This cone was identified as a blue cone by Behrens et al (2016). Arrow shows axon. **B.** Segmentation, e2006, blue cone pedicle (cone 2, dark blue) makes a variety of contacts with nearby rod spherules. 1. Telodendrial contacts close to the synaptic mouth of 4 rod spherules. **A.** 2. A roof contact, arrow, with a rod spherule (green). 3. Telodendrial contacs with a low rod (cyan) in the lowest row of the ONL. The contacts of blue cone pedicles are indistinguishable from green cones.

**Figure 5, supplement 5** Blue Cones e2006, 3D Reconstruction **A.** Reconstructed cone pedicles (cone 5, green and cone 2, dark blue, arrowhead, identified as a blue cone by Behrens et al (2016)). Arrows indicate ascending axons among ghost cells of the ONL (grey). **B.** The block of rods receiving contacts from 3 adjacent cone pedicles, cones 5 (green), cone 3 (cyan) and blue cone 2 (dark blue). **C.** Rotated view showing the top surface of these 3 cone pedicles with contact pads in red, yellow and orange. Arrows indicate ascending axons, arrowhead shows a ring-like structure of contacts. **D.** Rotated view, bottom surface of the rod spherules with contact pads, arrow head marks same ring of contacts, as in C. **E.** Detail, several rod spherules receiving contacts from the blue cone 2, orange, and cone 3, yellow. Rod spherules receive contacts indiscriminately from blue and green cones or both simultaneously. Arrows mark cone axons.

**Figure 7, supplement 1** Rod/rod and cone/cone contacts do not appear as gap junctions, FIB-SEM **A.** Partial reconstruction of 6 cone pedicles from FIB-SEM. **B.** Examples of cone/cone contacts (coral/blue or yellow/cyan, arrows), which show no membrane density. These examples were selected because of the nearby rod/cone contacts (arrowheads) which show prominent dense staining at their contact points, consistent with the presence of a gap junction. **C.** Rod/rod contacts (coral/magenta, arrows) show no membrane specialization, compared to nearby rod/cone gap junctions (coral/green, arrowheads).

## Videos

### Video 1 from **Fig. 1A**

Animation of a confocal series from the OPL showing Cx36 (red) decorating the cone telodendria (green, cone arrestin). All Cx36 clusters fall within the band of vGlut1 (blue) labeled rod spherules. No Cx36 in the ONL.

### Video 2 from **Fig. 1D**

3D reconstruction of a single EGFP labeled cone pedicle among all cone arrestin (blue) labeled cones, with extensive telodendria. All the Cx36 clusters in the OPL, then only those Cx36 clusters, more than 50, colocalized with the single EGFP labeled cone pedicle. Cx36 is distributed along the telodendria, often at the tips, and on the upper surface or roof of the cone pedicle.

### Video 3 from **Fig. 1**, supplement 2

Animating from the OPL to the outer segments following the cone axon allows cone labeled for blue cone opsin to be connected to its pedicle, labeled for cone arrestin.

### Video 4

3D reconstruction of a single cone (cone 5, green) with contact pads in red, then all rod spherules which were in contact with this cone.

### Video 5 from **Fig. 5F – K**

A pair of reconstructed adjacent cone pedicles (cone 5, green, & cone 3, cyan). A rotated view shows the contact pads from the top of the cone, then the contact pads from the bottom of the rod spherules, zooming in to show how a contact pad is close to the post-synaptic opening.

### Video 6 from **Fig. 6H**

3D reconstruction showing a large horseshoe-shaped gap junction (red) around the base of a transparent rod spherule with post-synaptic processes also filled (from FIB-SEM)

#### Source data

**Figure 1** – source data 1

Number of Cx36 clusters on a single cone pedicle.

Source data for Figure 1F.

**Figure 2** – source data 1

Number of Cx36 clusters on a rod spherule.

Source data for Figure 2F, G.

**Figure 3** – source data 1

Cone pedicle skeletons and rod spherule points.

Knossos data file of skeletons for Figure 3, 4.

**Figure 4** – source data 1

Cone coverage area and rod convergence.

Source data for Figure 4F-H.

**Figure 4** – source data 2

The number of rod spherules shared between two adjacent cone pairs.

Source data for Figure 4 – supplement 2B.

**Figure 7** – source data 1

Quantitative analysis of rod spherules from FIB-SEM

Source data for Figure 7A-G.

#### Data availability

All the data used to create the figures in the manuscript have been provided as source data files for Figures 1, 2, 3, 4, and 7.

The following data sets were generated. **In Process.**

Ishibashi M, Keung J, Ribelayga CP, Massey SC (2018) Confocal imaging of the outer plexiform layer in mouse retina. ID:xxx. URL:xxxx In the public domain at BIL http://www.brainimagelibrary.org/index.html

Singer JH (2018) SBF-SEM of mouse retina. eel001. URL:xxxx In the public domain at webKnossos https://webknossos.org/

Morgan CW, Aicher SA, Carroll JR (2019) FIB-SEM of the outer plexiform layer in light-adapted mouse retina. EM1 and EM2, ID:xxx. URL:xxxx In the public domain at BossDB https://bossdb.org/

## Contributions

The project was designed by SCM, MI, IF and CPR; JS contributed an SBF-SEM dataset; CWM, SAA and JRC prepared two FIB-SEM datasets; LJ and WL contributed the blue cone mouse line; MI and SCM performed confocal microscopy and data analysis; JK performed immunocytochemistry; SCM, MI, IF and CPR, first draft and editing; all authors edited and approved final draft.

## Acknowledgement

This project was inspired by the paper from Behrens et al (2016) who used e2006 to reconstruct bipolar cells. We thank Christian Behrens, Timm Schubert, Philipp Berens and Thomas Euler (University of Tübingen) for generously sharing data on blue cone bipolar cells. We thank Moritz Helmstaedter (MPI, Frankfurt) for hosting the e2006 dataset. We thank David Berson, Brown University, for advice, encouragement and an introduction to connectomics. We thank Jessica Riesterer at the Multiscale Microscopy Core, an OHSU University Shared Resource core facility, for acquiring the FIB-SEM datasets. Supported by NIH grants: EY017836 (JHS); EY02948 (SCM & CPR); P30EY028102 (SCM); P30NS061800 (SAA); R01MH127343 (SAA, SCM, CWM, & CPR)

