## Supplementary figures and images for "Analysis of Rod/Cone Gap Junctions from the Reconstruction of Mouse Photoreceptor Terminals"

### Figure 1, supplement 1

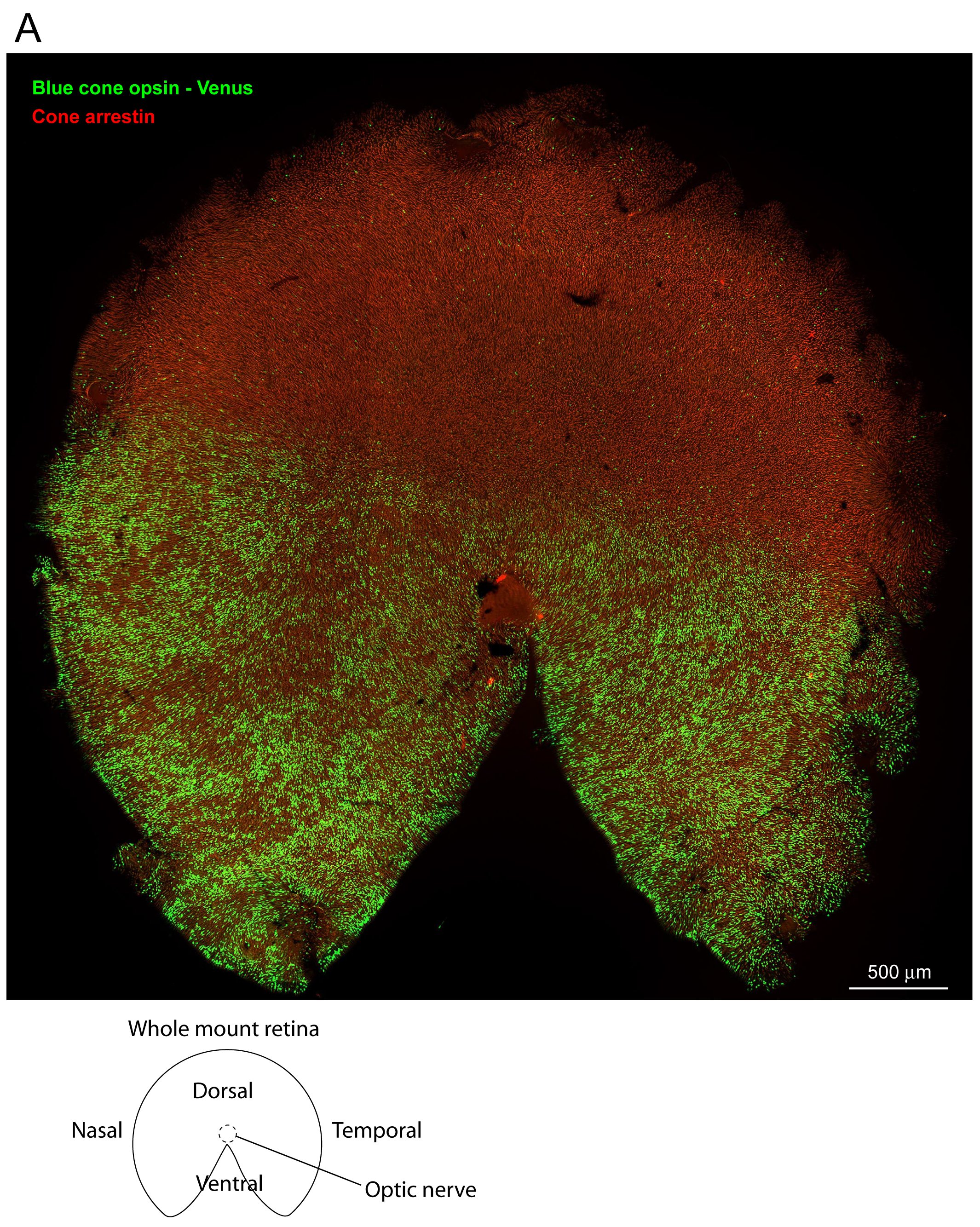

### Figure 1, supplement 2

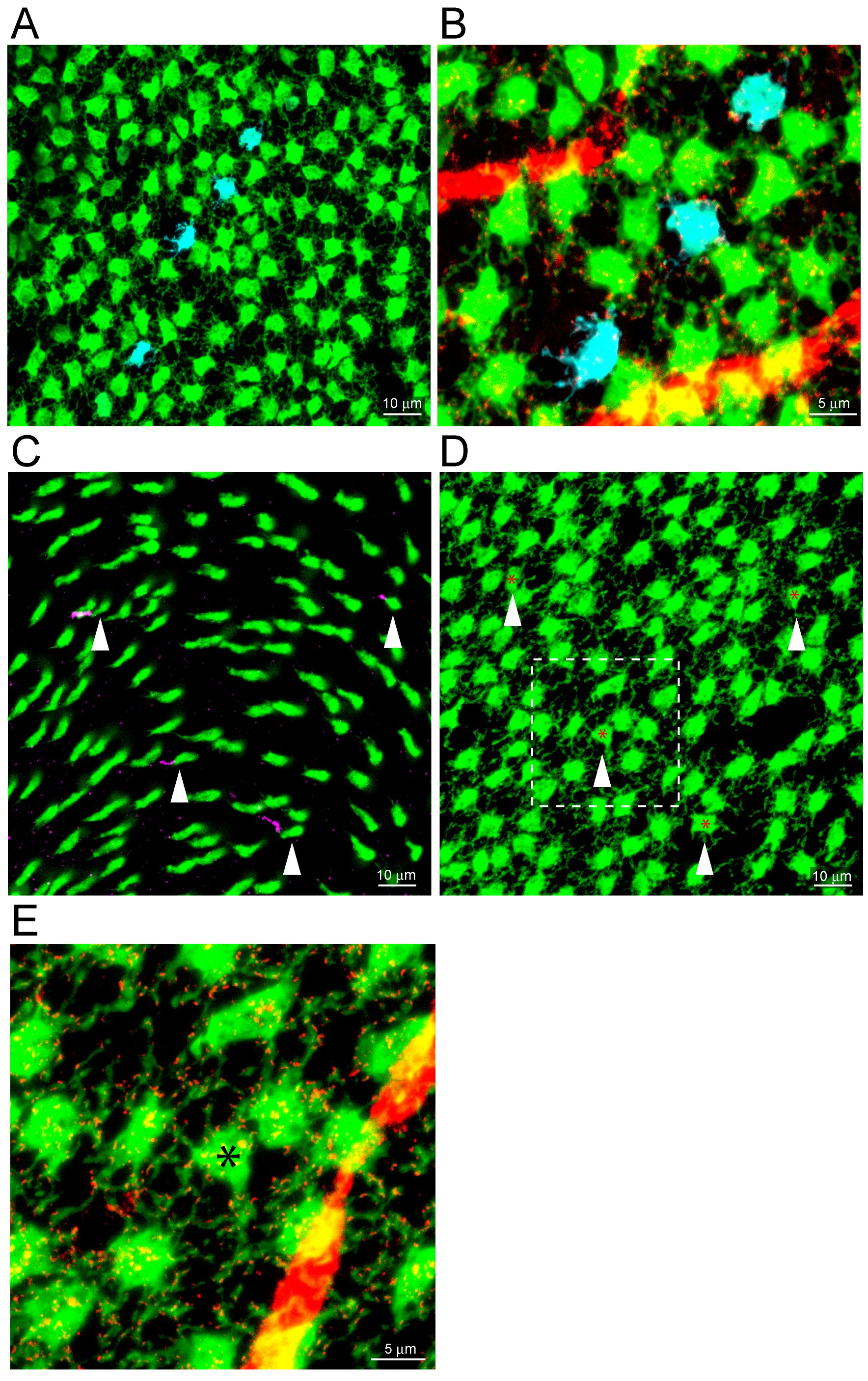

### Figure 4, supplement 1

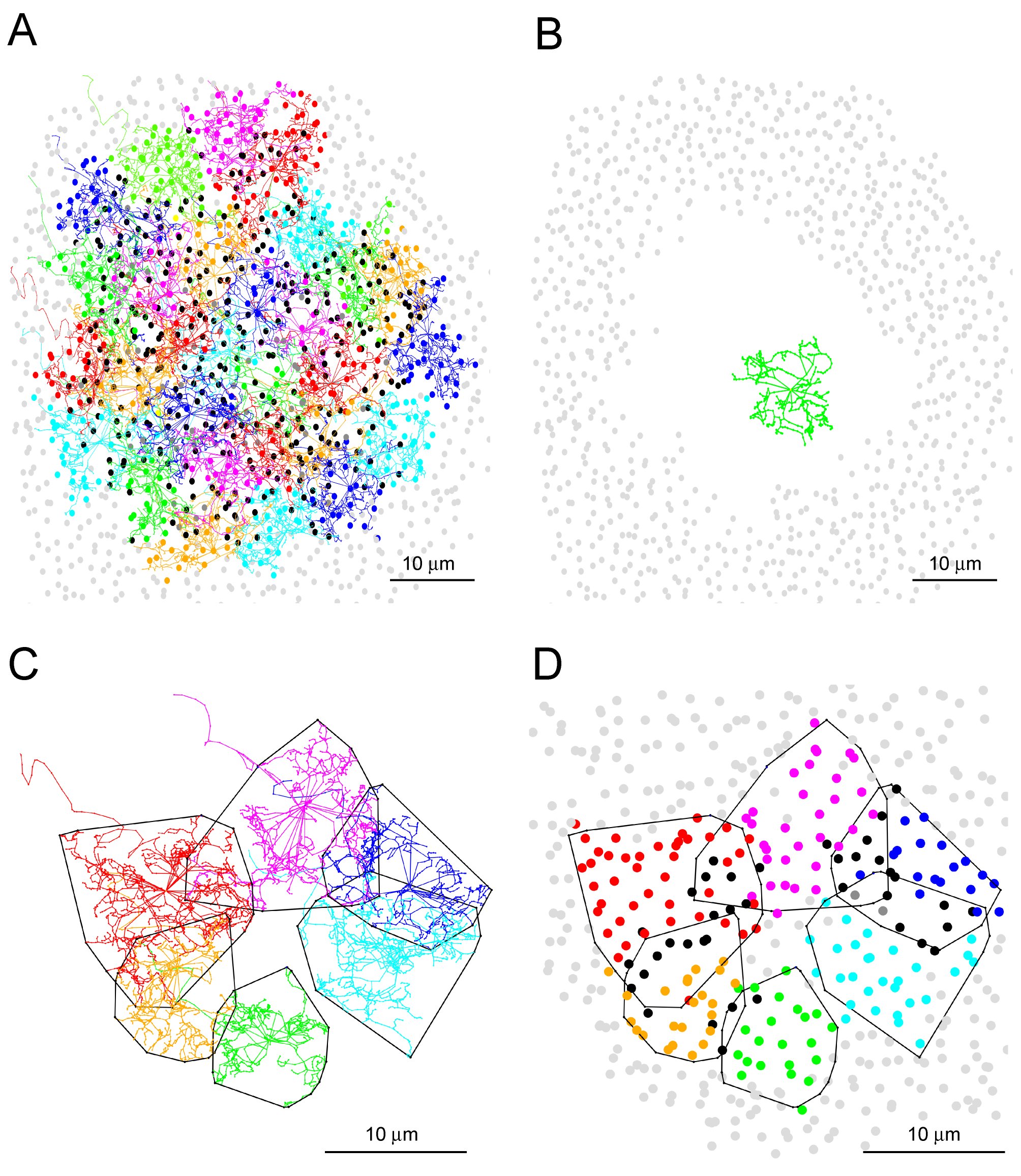

### Figure 4, supplement 2

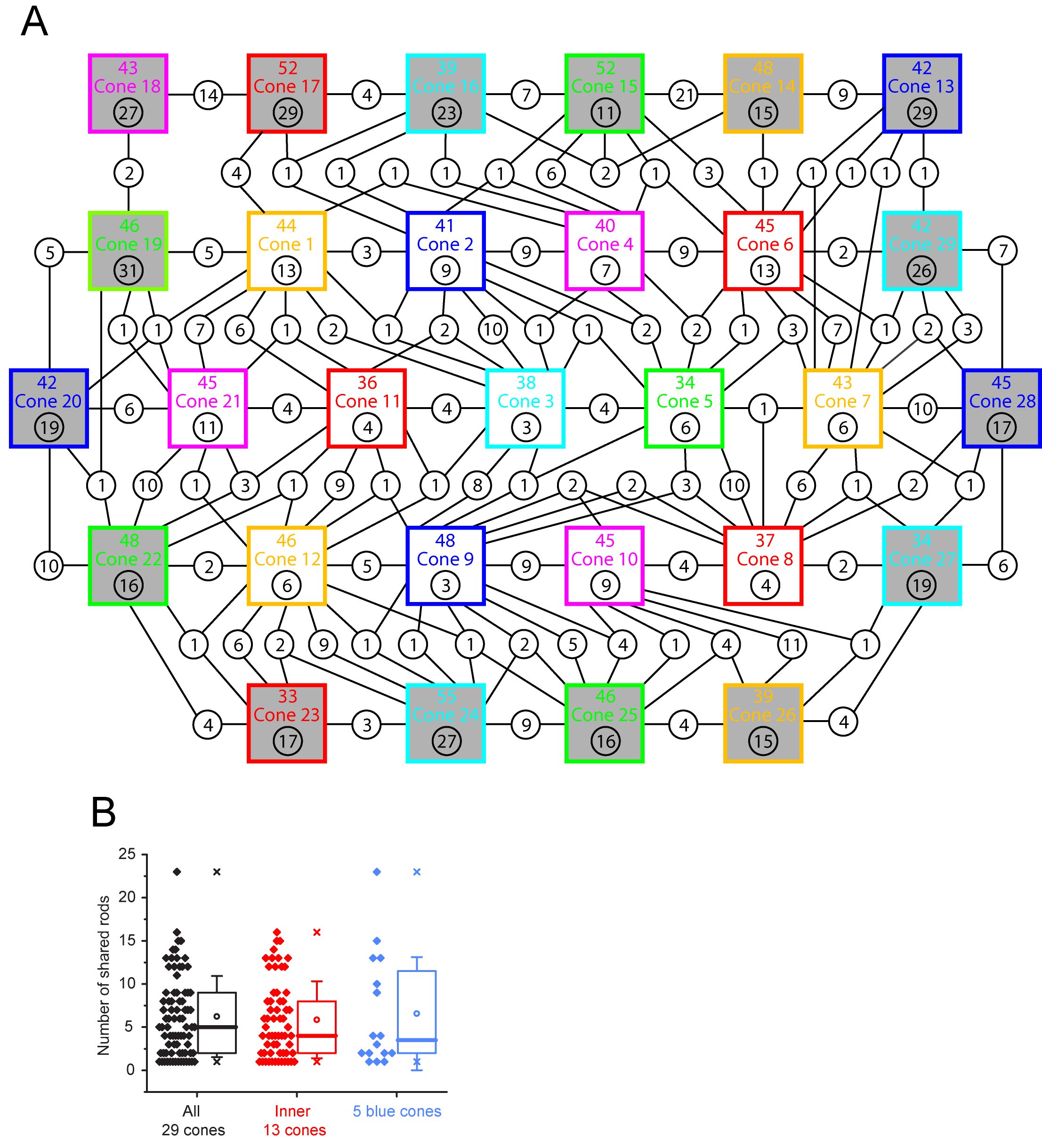

### Figure 5, supplement 1

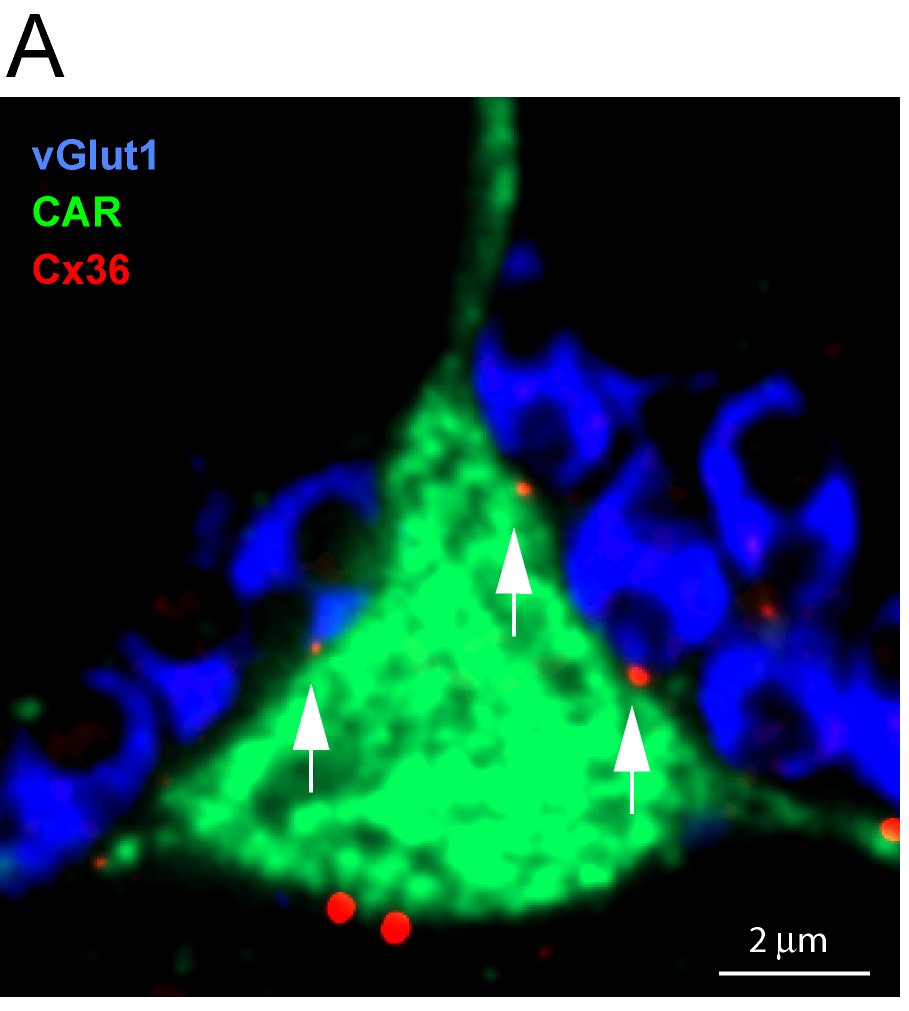

### Figure 5, supplement 2

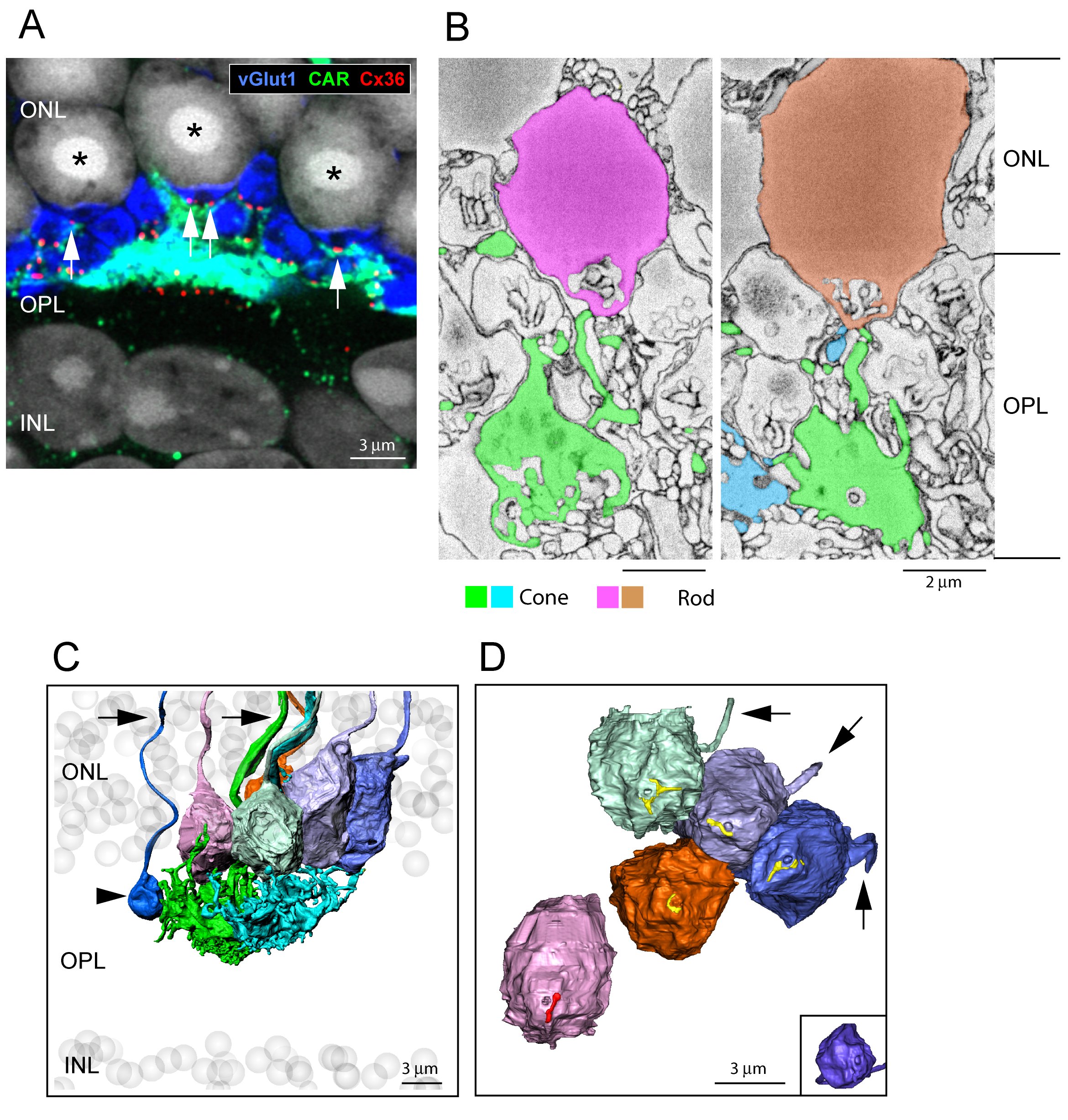

### Figure 5, supplement 3

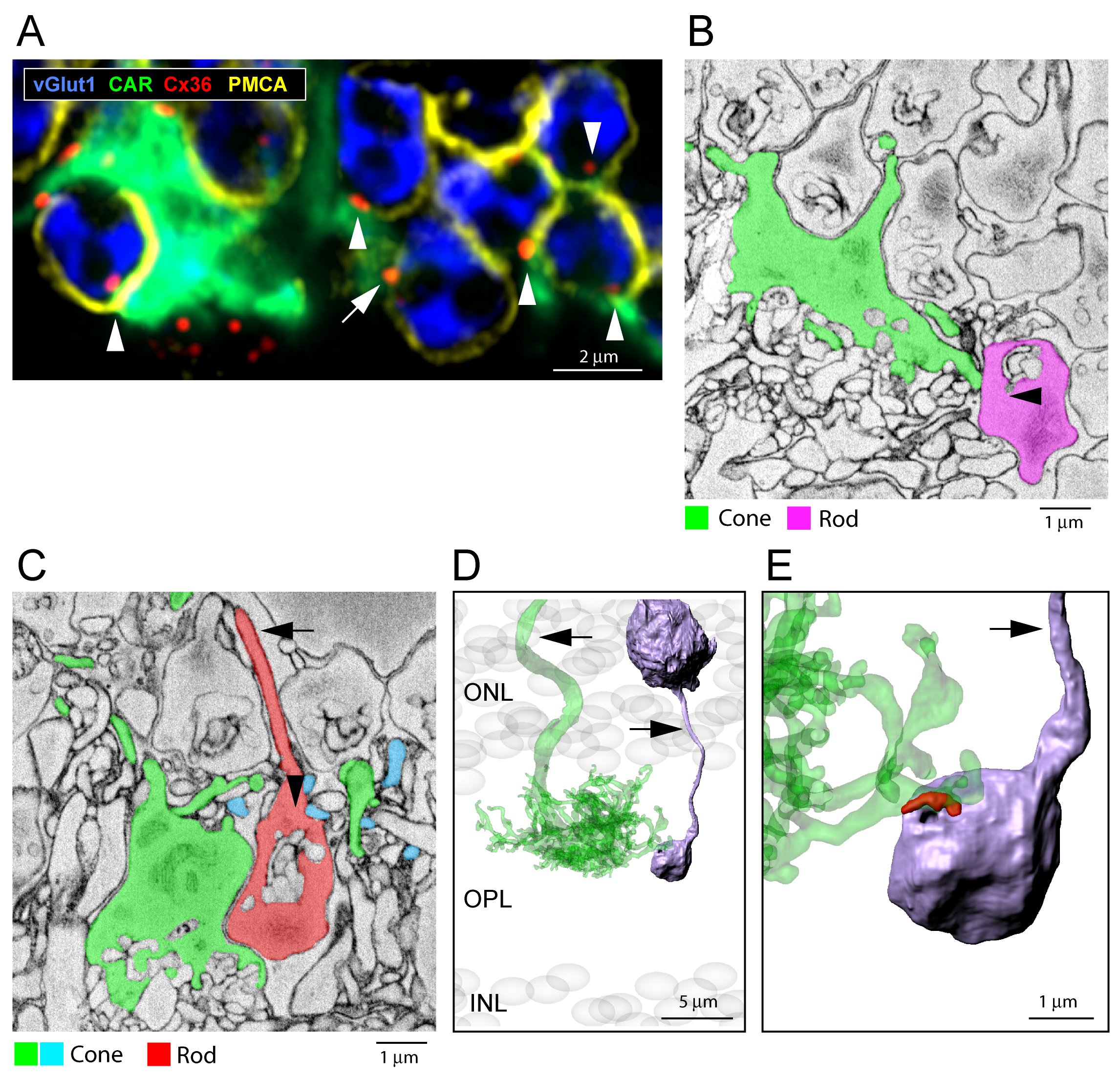

### Figure 5, supplement 4

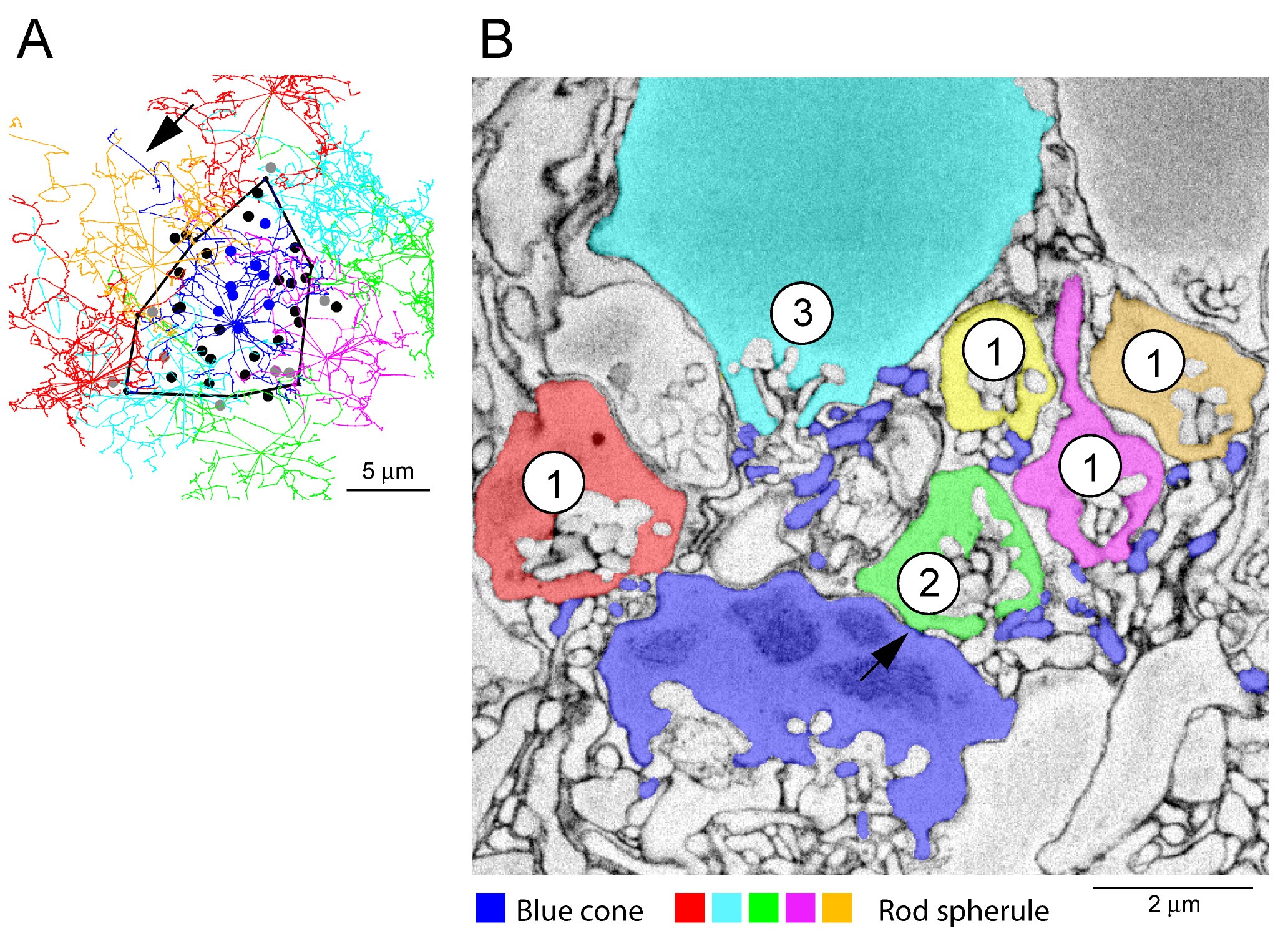

### Figure 5, supplement 5

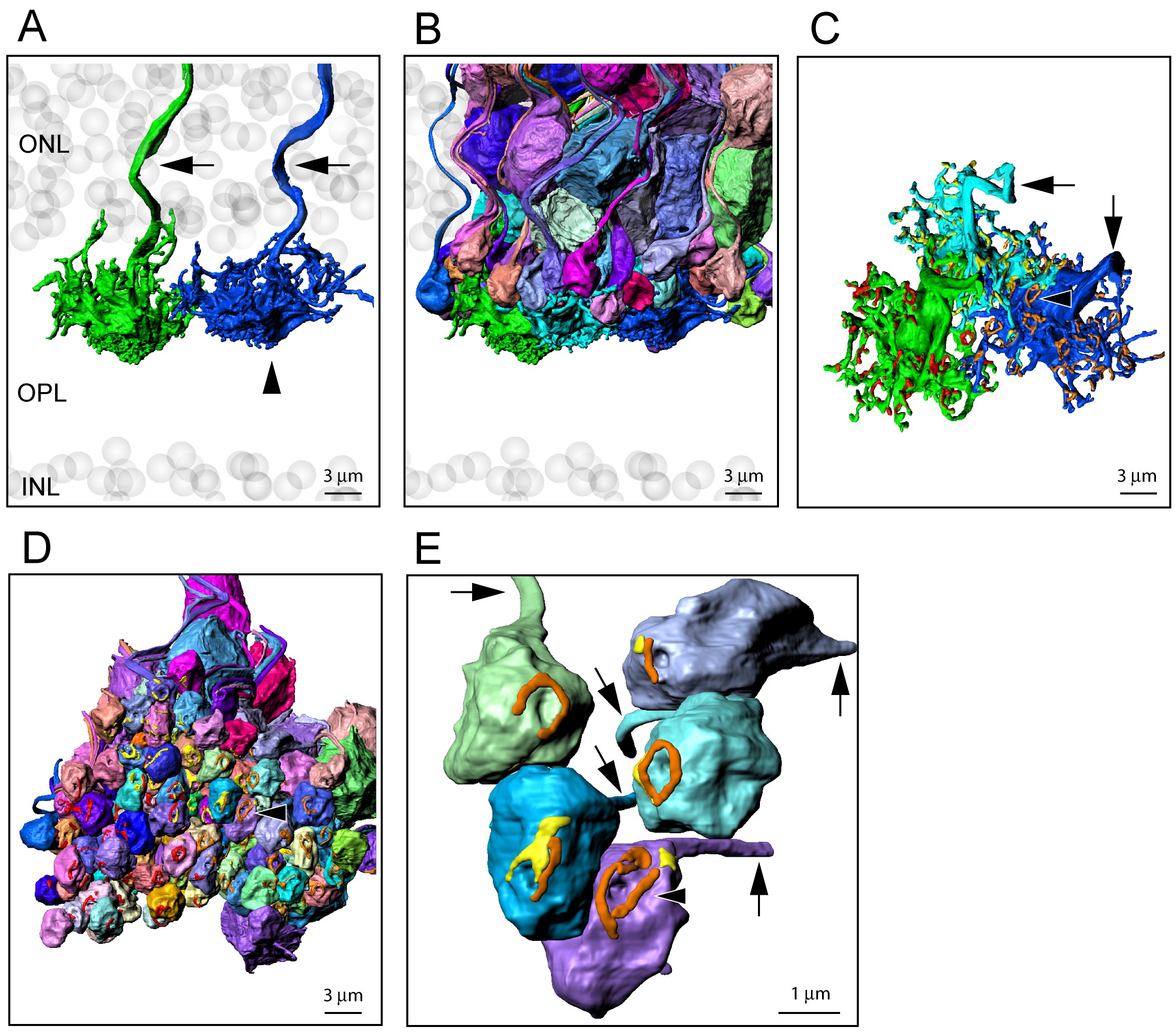

### Figure 7, supplement 1

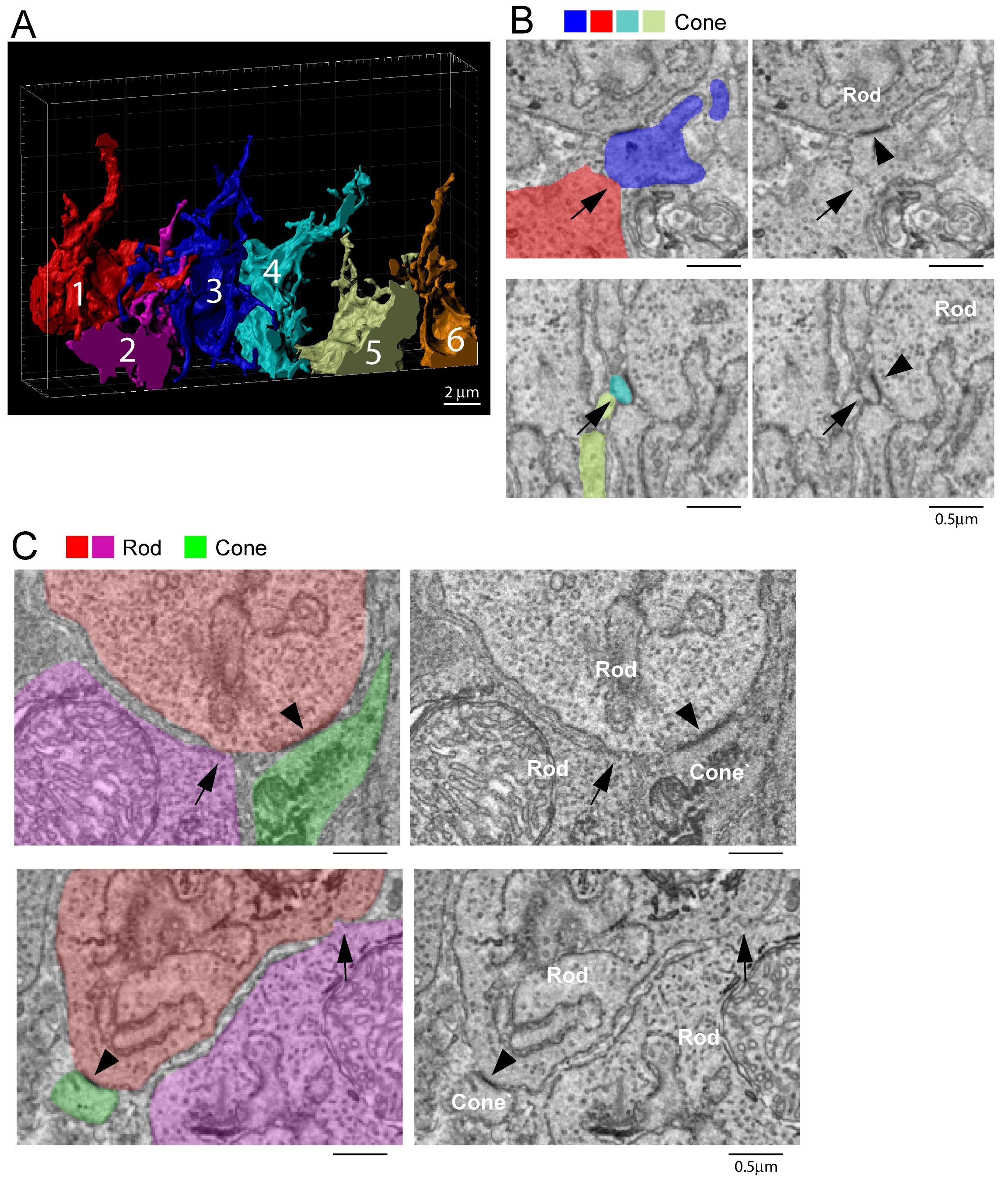
